# A conserved long intergenic non-coding RNA containing snoRNA sequences, *lncCOBRA1*, affects Arabidopsis germination and development

**DOI:** 10.1101/2021.11.28.470209

**Authors:** Marianne C. Kramer, Hee Jong Kim, Kyle R. Palos, Benjamin A. Garcia, Eric Lyons, Mark A. Beilstein, Andrew D.L. Nelson, Brian D. Gregory

## Abstract

Long non-coding RNAs (lncRNAs) are an increasingly studied group of non-protein-coding transcripts with a wide variety of molecular functions gaining attention for their roles in numerous biological processes. Nearly 6,000 lncRNAs have been identified in *Arabidopsis thaliana* but many have yet to be studied. Here, we examine a class of previously uncharacterized lncRNAs termed *<u>CO</u>NSERVED IN* <u>B</u>RASSICA <u>RA</u>PA (*lncCOBRA*) transcripts that were previously identified for their high level of sequence conservation in the related crop species *Brassica rapa*, their nuclear-localization and protein-bound nature. In particular, we focus on *lncCOBRA1* and demonstrate that its abundance is highly tissue and developmental specific, with particularly high levels early in germination. *lncCOBRA1* contains two snoRNAs domains within it, making it the first sno-lincRNA example in a non-mammalian system, though we find that it is processed differently than its mammalian counterparts. We further show that plants lacking *lncCOBRA1* display patterns of delayed gemination and are overall smaller than wild-type plants. Lastly, we identify the proteins that interact with *lncCOBRA1* and examine the protein-protein interaction network of *lncCOBRA1*-interacting proteins.

## INTRODUCTION

Long non-coding RNAs (lncRNAs) are transcripts defined as greater than 200 nucleotides (nt) that lack or have an open reading frame less than 100 amino acids (Liu et al., 2012). Transcriptome-wide studies have demonstrated that lncRNAs are often expressed in a context-specific manner, a characteristic believed to facilitate some of their known functions in modulating gene expression, mRNA splicing, and translation (Quinn and Chang, 2016). The function of lncRNAs is highly dependent on their subcellular location. Nuclear lncRNAs often serve key roles in regulating gene expression, either in *cis* (neighboring genes) or in *trans* (distant genes). LncRNAs can also bind and sequester proteins, such as proteins involved in chromatin stability and splicing factors, from their target chromosomal regions, thereby affecting gene expression (Lee et al., 2016; Yin et al., 2012).

In plants, lncRNAs are implicated in numerous biological mechanisms with demonstrated functions in flowering, organogenesis, photomorphogenesis, reproduction, and abiotic/biotic stress responses (reviewed in Wang and Chekanova, 2017). Most research has focused on the intergenic class of lncRNAs (lincRNAs) (Mattick and Rinn, 2015), as historically it has been easier to discern their transcriptional origins relative to other lncRNAs that overlap protein-coding genes (e.g., natural antisense transcripts (NATs)). In plants, detailed annotation and functional efforts have led to the identification of several lincRNAs with characterized functions in regulation of auxin signaling outputs (Ariel et al., 2014), response to drought/salt stress and resistance to pathogens (Seo et al., 2017; Qin et al., 2017), and response to phosphate starvation (Franco-Zorrilla et al., 2007).

While most Pol II transcribed lincRNAs are 5’ capped and 3’ polyadenylated, recently a previously uncharacterized group of lncRNAs that lacks one or both of these features has been described (Xing and Chen, 2018). These non-canonical Pol II-dependent lncRNAs have small nucleolar RNA (snoRNA) sequences at their 5’ and 3’ ends and are referred to as sno-lncRNAs. snoRNAs are 70 – 200 nt highly structured, nuclear-localized, protein-bound non-coding RNAs that are usually concentrated in the Cajal bodies or nucleolus (Reichow et al., 2007). SnoRNAs co-transcriptionally form snoRNA-ribonucleoprotein complexes (snoRNPs) (Kiss, 2001) and function through complementarity with ribosomal RNA (rRNA) sequences to guide rRNA modification to ultimately participate in ribosome subunit maturation. In one example, the formation of snoRNPs at the ends of sno-lncRNAs protects the intervening sequence from exonuclease trimming (Yin et al., 2012).

sno-lncRNAs have been identified in humans, rhesus monkeys, and mice (Yin et al., 2012; Zhang et al., 2014; Xing et al., 2017) but have yet to be described in plants. A functional analysis of sno-lncRNAs in humans was recently performed, where SLERT was identified (snoRNA-ended lncRNA enhances pre-ribosomal RNA transcription; (Xing et al., 2017)). SLERT localizes to the nucleolus in a manner dependent on the two snoRNPs at its ends and functions to promote active transcription of rRNAs (Xing et al., 2017). Thus, sno-lncRNAs represent an interesting class of lncRNAs with evident functions in humans.

Due to their lack of protein-coding capacity, lincRNAs typically display poor sequence conservation among even closely related species (Nelson et al., 2016; Necsulea et al., 2014). LincRNAs with functions defined by structural or sequence-specific interactions with other molecules (e.g., proteins) will likely display higher levels of conservation over lincRNAs that function based on proximity to other genes (e.g., transcription enhancers/repressors). We previously identified lincRNAs in the nuclei from 10-day-old seedlings and found that lincRNAs with RNA binding protein (RBP) binding sites were significantly more likely to be conserved at the sequence-level in *Brassica rapa* than those that lacked protein binding sites (Gosai et al., 2015), suggesting these protein-bound, conserved lincRNAs may be of functional importance in plants.

Here, we assess the function of those nuclear, protein-bound, and conserved lincRNAs that we have termed *CONSERVED IN* BRASSICA RAPA (*lncCOBRA*). We find that the *COBRA* lincRNAs display germination- and developmental-dependent patterns of abundance and, in particular, we focus on *lncCOBRA1* which contains two snoRNA sequences within it, indicating the first evidence of a sno-lincRNA in Arabidopsis. Unlike sno-lncRNAs identified in humans, *lncCOBRA1* is transcribed from an intergenic region, and is transcribed as a longer transcript before processing at its 3’ end. We further show that *lncCOBRA1* influences plant germination and growth, as plants lacking *lncCOBRA1* germinate later and are smaller than wild type plants. Lastly, we identify *lncCOBRA1*-interacting proteins, including the scaffold protein RACK1A, and several of its known interactors.

## RESULTS

### Identification of conserved, nuclear, protein-bound long intergenic non-coding RNAs (lincRNAs)

We previously identified 236 nuclear lincRNAs from 10-day-old seedlings, of which 38 contained up to four RNA-binding protein (RBP) interaction sites (Gosai et al., 2015). These protein-bound lincRNAs were significantly more conserved within the related crop species *Brassica rapa* than those lacking RBP binding sites (**Supplemental Figure 1A** and **Table 1**) (Gosai et al., 2015). Since lincRNAs do not encode proteins, small polymorphisms within the sequence generally have little functional consequence, and thus lincRNAs are generally not well conserved at the sequence level (Hezroni et al., 2015; Necsulea et al., 2014; Ponjavic et al., 2007). Thus, the combination of conservation in *Brassica rapa* and nuclear protein binding suggested that these lincRNAs may have important functions in plant systems and were named *<u>CO</u>NSERVED IN <u>B</u>RASSICA <u>RA</u>PA 1-14* (*lncCOBRA1-14*) (**Supplemental Figure 1A** and **Table 1**).

We initiated our search for function by examining their abundance during seed germination, as lincRNAs in several eukaryotic species are essential during development (e.g. *HOTAIR*, *COOLAIR*) (Liu et al., 2012; Swiezewski et al., 2009; Rinn et al., 2007; Sarropoulos et al., 2019). Using a previously published transcriptomic dataset (Narsai et al., 2017), we found that the majority (N = 9; 64%) of *lncCOBRA* transcripts displayed germination-dependent patterns of abundance, with peaks in abundance at various points during seed germination (**Supplemental Figure 1B**). Going forward, we focused on *lncCOBRA1*, *lncCOBRA3*, and *lncCOBRA5* due to their highly specific abundance profiles during seed germination and the availability of insertional mutant lines for these loci. *lncCOBRA1* and *lncCOBRA3* were most abundant after 48 hours of stratification at 4°C in the dark followed by one hour in light, while *lncCOBRA5* abundance was highest slightly later, with a peak in abundance six hours after transfer into light conditions (**Figure 1A** and **Supplemental Figure 1B**). Abundance of the three *lncCOBRA* transcripts decreased rapidly as the seeds progressed through germination and transitioned into seedlings (**Figure 1A** and **Supplemental Figure 1B**). Supporting this, the Arabidopsis expression atlas in the eFP Browser (Klepikova et al., 2016) revealed that all three *lncCOBRA* transcripts were expressed early during seed germination, with the highest expression at one hour after imbibition (**Supplemental Figure 1C**). The abundance of *lncCOBRA1*, *lncCOBRA3*, and *lncCOBRA5* was also dynamic throughout seedling development as measured by quantitative reverse transcription-PCR (qRT-PCR), as they had the highest abundance in 2-day-old seedlings and rapidly decreased in abundance as the seedlings aged (**Figure 1B** and **Supplemental Figure 1D**).

**Figure 1:**
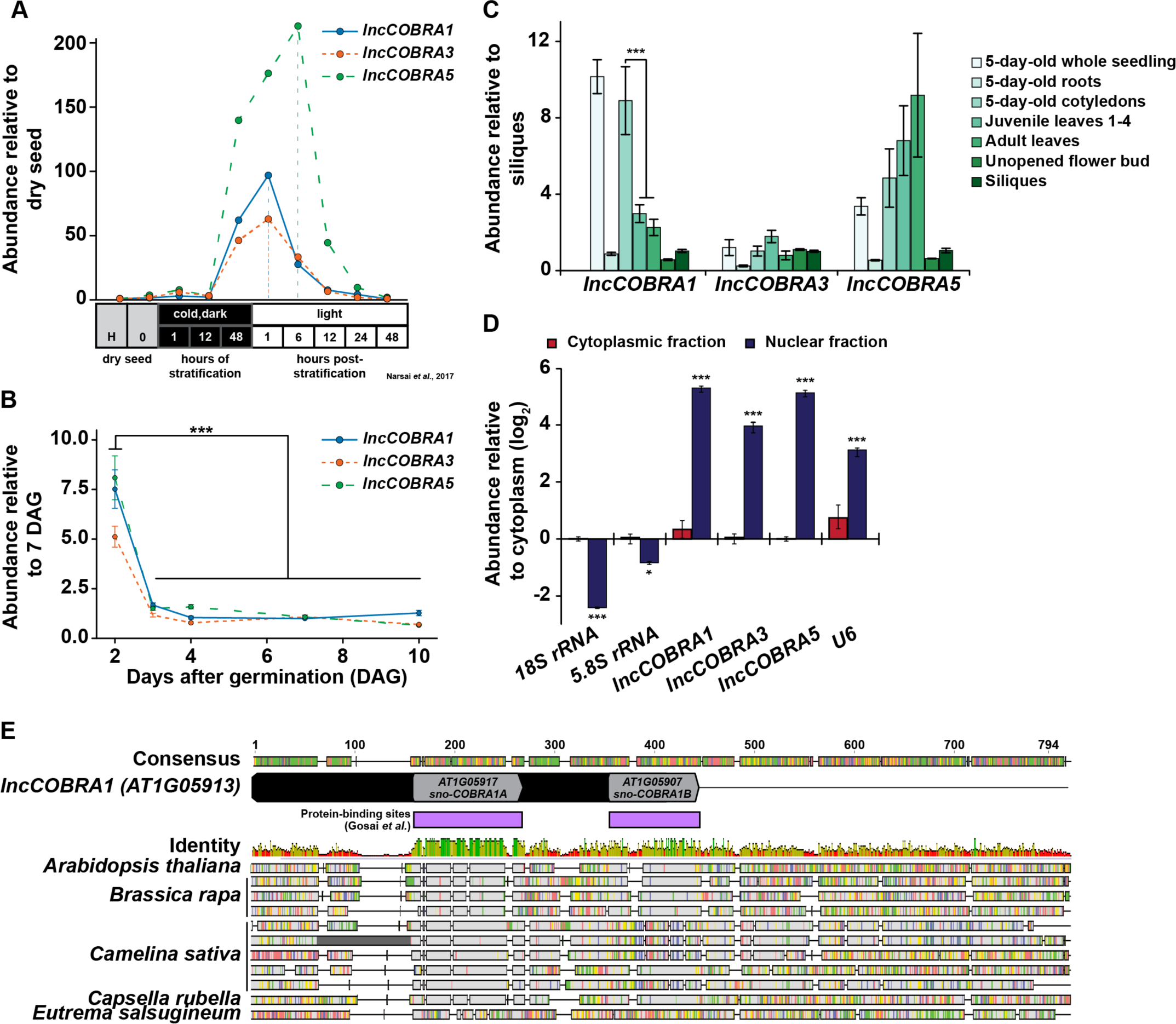
Identification and expression of highly conserved, protein-bound lincRNA, lncCOBRA1. (A) Abundance of *lncCOBRA1*, *lncCOBRA3*, and *lncCOBRA5* during germination as previously measured by RNA-seq (Narsai et al., 2017). Abundance is relative to dry seed after harvest. (B) Abundance of *lncCOBRA1*, *lncCOBRA3*, and *lncCOBRA5* during early seedling development as measured by qRT-PCR. Abundance is normalized by the geomean of *UBC9* and *UBC10* and relative to 7-day-old seedlings. *** denotes p-value < 0.001, Wilcoxon t-test. (C) Abundance of *lncCOBRA1*, *lncCOBRA3*, and *lncCOBRA5* in various tissues as measured by qRT-PCR. Abundance is normalized by the geomean of *UBC9* and *UBC10* and is relative to siliques seedlings. *** denotes p-value < 0.001, Wilcoxon t-test. (D) Abundance of *lncCOBRA1*, *lncCOBRA3*, and *lncCOBRA5* in nuclear and cytoplasmic fractions as measured by qRT-PCR. Abundance is normalized to *UBC9* and relative to cytoplasmic fraction. *18S rRNA* and *5.8S rRNA* are cytoplasmic positive controls and *U6* is a nuclear positive control. *, *** denotes p-value < 0.05, < 0.001, Wilcoxon t-test. (E) Conservation *lncCOBRA1* in *Brassica rapa*, *Camelina sativa*, *Capsella rubella*, and *Eutrema salsugineum.* Conservation was examined using Geneious Prime (Geneious | Bioinformatics Solutions for the Analysis of Molecular Sequence Data, 2019). Protein-binding sites were identified in the nuclei from 10-day-old seedlings in (Gosai et al., 2015). Colors in identity: Green = 100%, green-brown = 30-100%, red < 30% identity.

Additionally, *lncCOBRA* transcripts displayed tissue specific patterns of accumulation. We found that *lncCOBRA5* abundance was highest in leaf tissue and increases in abundance as the age of the leaf progressed from embryonic cotyledons to juvenile leaves and adult leaves (**Figure 1C**). In contrast, *lncCOBRA1* had the highest abundance in 5-day-old seedlings, specifically in the cotyledons, and its abundance decreased as the leaves increased in age, with a significant (p-value < 0.001; Wilcoxon *t* test) decrease in abundance between 5-day-old cotyledons and true leaves (both juvenile and adult leaves) (**Figure 1C** and **Supplemental Figure 1E**). Thus, all three *lncCOBRA* transcripts examined were highly abundant early in germination and decreased as development progressed. In particular, *lncCOBRA1* was highly abundant in embryonic cotyledons and decreased in abundance as true leaves emerge, suggesting *lncCOBRA1* may function during germination and/or early in plant development.

Since these lincRNAs were originally identified as nuclear lincRNAs, and lincRNA function is influenced by subcellular localization, we sought to determine if they were nuclear retained. To do so, we isolated pure nuclear and cytoplasmic fractions using the isolation of nuclei tagged in specific cell types (INTACT) technique (Deal and Henikoff, 2010, 2011) and performed qRT-PCR for *lncCOBRA1*, *lncCOBRA3*, and *lncCOBRA5* as well as nuclear (*U6*) and cytoplasmic (*5.8S rRNA* and *18S rRNA*) positive controls. All three *lncCOBRA* transcripts were significantly (p-value < 0.001; Wilcoxon t-test) enriched in the nuclear fraction, like *U6*, compared to the cytoplasmic fraction where the two rRNAs were enriched, confirming these transcripts were indeed primarily nuclear localized (**Figure 1D**).

### *lncCOBRA1* contains two highly-conserved snoRNA domains and is processed at its 3’ end after transcription

As both *lncCOBRA1* and *lncCOBRA3* contain small nucleolar RNA (snoRNA) sequences annotated within their transcripts (**Figure 1E** **and Supplemental Figure 1F**), and given the evident importance of sno-lncRNAs in humans (Xing and Chen, 2018), we were particularly interested in these two transcripts. Since *lncCOBRA3* lacked tissue-specific patterns of abundance and *lncCOBRA1* had distinct patterns of abundance during seed germination and development, we decided to focus on *lncCOBRA1*. *lncCOBRA1* was annotated to be a 318 nt lincRNA in the Araport11 genome annotation and contained two snoRNA sequence domains within it. The two annotated snoRNA domains overlapped with two previously identified RBP binding sites (**Figure 1E**) (Gosai et al., 2015). These RNA binding/snoRNA domains displayed the highest level of sequence similarity in a sequence alignment of *lncCOBRA1* homologs from five Brassicaceae with *AT1G05917* (sno-*COBRA1A)* and *AT1G05907* (sno-*COBRA1B)* having ∼79% and 56% sequence identity among the profiled species, respectively (**Figure 1E**). *lncCOBRA1* was highly conserved in all species profiled, with 30-46% sequence identity in the 500 nt up- and downstream of the 5’ most snoRNA (*AT1G05917*), which also included *sno-COBRA1B* (**Figure 1E** and **Supplemental Figure 2A**) (Geneious | Bioinformatics Solutions for the Analysis of Molecular Sequence Data, 2019). To ensure we are examining the *lncCOBRA1* lincRNA rather than a functional set of snoRNAs, two primer sets were used for all qRT-PCR analyses, one set within sno-*COBRA1A* and the other set (set 2) amplifying the region between the two snoRNAs (**Supplemental Figure 1G;** blue and red primers). In addition to their sequence conservation within Brassicaceae, sno-*COBRA1A* and sno-*COBRA1B* have sequence homology to human SNORD59A and SNORD59B, with sequence identity of 76% and 90%, respectively (Liang-Hu et al., 2001) (**Supplemental Figure 2B**). In fact, their tandem orientation is also conserved in humans, with SNORD59A upstream of SNORD59B in an intron of the protein-coding transcript encoding ATP synthase subunit d (ATP5PD) (Kiss-László et al., 1996). Overall, these findings indicate that these snoRNA sequences and orientation are highly conserved, suggesting they are of significant evolutionary importance.

In humans, sno-lncRNAs are derived from introns excised from protein-coding mRNAs that contain two snoRNA sequences (Xing and Chen, 2018). Instead of being degraded like normal, these introns are debranched and trimmed at the 5’ and 3’ ends by exonucleases until the enzyme reaches the snoRNA domain. The highly structured and protein-bound nature of the snoRNA sequences acts as protection from further degradation, resulting in lncRNAs flanked by snoRNA sequences at each end, but that lack 5’ caps and poly(A) tails (Xing and Chen, 2018). To determine if a similar mechanism was used during *lncCOBRA1* biogenesis, we performed 5’ rapid amplification of cDNA ends (5’ RACE) to determine the 5’ end of the transcript. In 5’ RACE, 5’ caps are removed, and an adapter is directly ligated to the 5’ end of RNA. Following reverse transcription with a gene specific primer and two rounds of PCR, the precise 5’ end of the transcript can be determined (**Figure 2A**). If the 5’ end of *lncCOBRA1* was as annotated in Araport11, we would expect PCR products of 250 and 319 bp produced with a primer within the 5’ adapter and two reverse primers, A and B, respectively (**Figures 2A-B**; red arrows). Indeed, the 5’ RACE PCR reactions produced products as expected, indicating that the annotated 5’ end of *lncCOBRA1* is indeed where the transcript begins (**Figures 2A-B**; red triangle), and thus *lncCOBRA1* is apparently not trimmed at the 5’ end after transcription.

**Figure 2:**
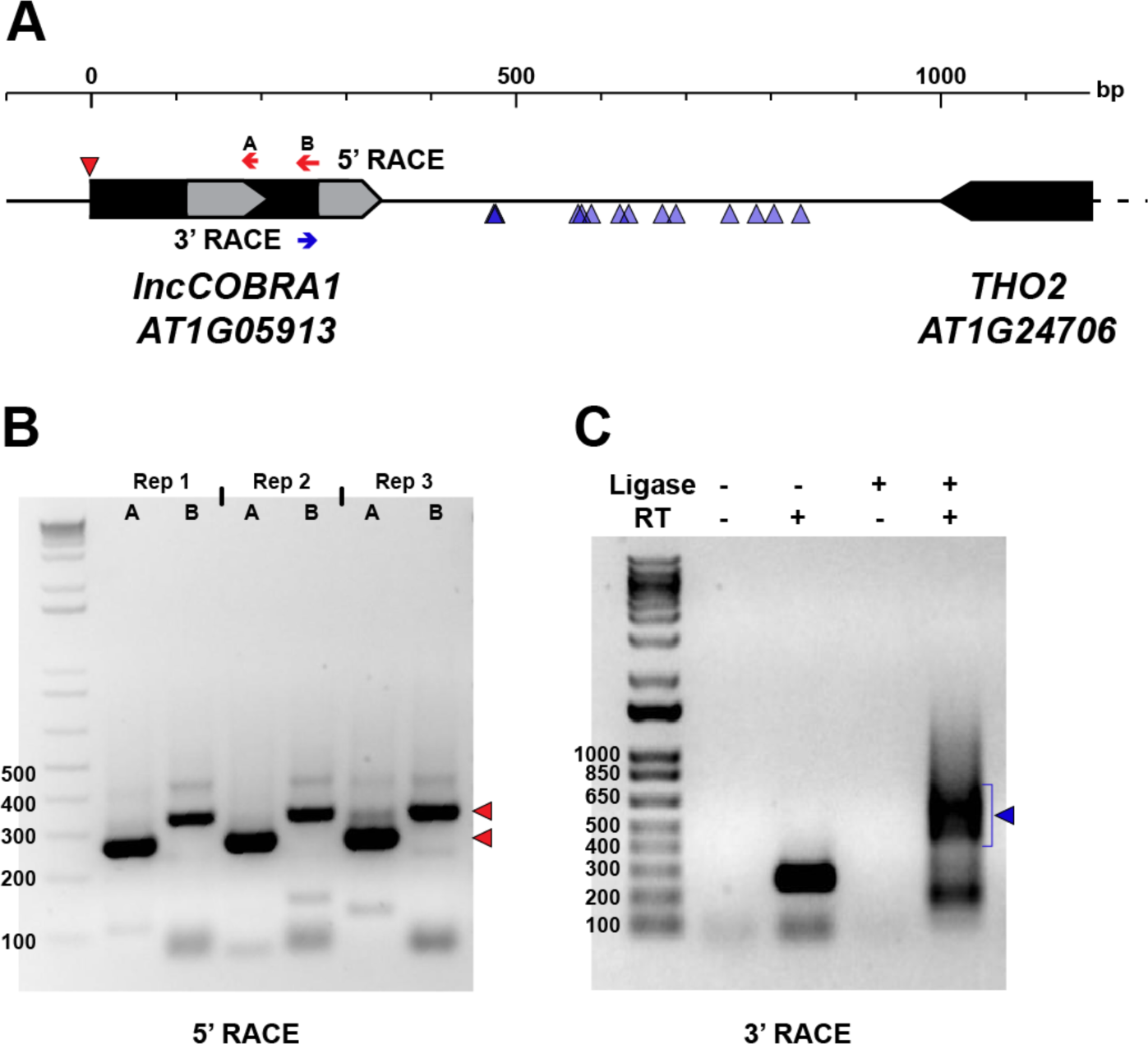
Post-transcriptional processing of *lncCOBRA1*. (A) Diagram of *lncCOBRA1* (*AT1G05913*) locus. Gray arrows represent the two snoRNAs annotated within *lncCOBRA1.* Red arrows represent the two primers used for 5’ RACE and red triangle represents the 5’ end identified by 5’ RACE PCR in Figure 2B. Blue arrow represents the primer used for 3’ RACE. Blue triangles represent the 3’ most end identified through Sanger sequencing 14 colonies. (B) Three biological replicates of 5’ RACE with primers indicated in Figure 2B. Red triangles represent the two major bands of PCR product. Ladder is 1 kb+. (C) PCR results from 3’ RACE in Col-0 5-day-old seedlings. -/+ T4 RNA ligase, -/+ SuperScript II. Ladder is 1 kb+.

We next asked if there was 3’ end processing and sought to determine the full length of *lncCOBRA1.* To begin, we performed RT-PCR with a forward primer at the 5’ most end of the transcribed RNA as confirmed by 5’ RACE and five tiled reverse primers (**Supplemental Figure 3A,** green arrows). This revealed that *lncCOBRA1* was substantially longer than originally annotated, with amplification of *lncCOBRA1* with all reverse primers, indicating that *lncCOBRA1* is transcribed as a much longer transcript, possibly over 1000 nt long (**Supplemental Figure 3B**). Given the tissue specificity of *lncCOBRA1* abundance (**Figure 1**), we performed the RT-PCR in 2-, 3-, 4-, and 5-day-old seedlings as well as seeds 1- and 2-days-after-imbibition to determine if there were different isoforms in a developmental manner. This revealed amplification with all reverse primers in all developmental time points, revealing that *lncCOBRA1* was over 1000 nt at these stages as well (**Supplemental Figure 3B**). Overall, this suggests that *lncCOBRA1* is a much longer lincRNA than initially hypothesized.

To determine the precise 3’ end of *lncCOBRA1*, we performed 3’ RACE. Similar to 5’ RACE, an adapter is ligated to the 3’ end followed by reverse transcription with a gene specific primer and two rounds of nested PCR (**Figure 2A**). The final PCR reaction produced a diffuse band around 500-650 bp in length, which would suggest a 742-892 nt long transcript based on the site of the 3’ RACE primer (**Figure 2A**, blue arrow; **Figure 2C**). Since the resulting 3’ RACE PCR band was diffuse, we extracted the PCR product, cloned it into a sequencing vector and performed Sanger sequencing to identify the precise 3’ end of *lncCOBRA1*. After sequencing 14 independent colonies, several 3’ ends of *lncCOBRA1* were revealed, with the majority of 3’ ends centering ∼250 and ∼350 nt downstream of the 3’ RACE primer (**Figure 2A**; blue triangles). The various 3’ ends detected by 3’ RACE, the diffuse 3’ RACE PCR band (**Figure 2C**), and the RT-PCR results (**Supplemental Figure 3B**) indicate that *lncCOBRA1* is transcribed as a longer transcript, possibly over 1000 nt in length (**Supplemental Figure 3B**) and is trimmed from its 3’ end to reach a final transcript ∼500-600 nt long, possibly with several stable 3’ ends. Importantly, in all of the 14 colonies sequenced, no polyA tail was identified. This, along with our inability to detect *lncCOBRA1* in any published polyA-selected RNA-seq datasets (data not shown) suggests that *lncCOBRA1* is not polyadenylated in its final processed form.

In plants, polycistronic snoRNAs are encoded in intergenic regions, transcribed by RNA Pol II and generally contain two conserved promoter elements, a *Telo*- box and a Site II element (combined referred to as *Telo*SII) (Gaspin et al., 2010). Notably, in Arabidopsis nearly all ribosomal protein genes and other genes involved in ribosome biogenesis and translation contain *Telo*SII elements in their promoters (Gaspin et al., 2010). This combined *Telo*SII element is found upstream of the TATA box and acts to coordinate expression of snoRNAs and protein-coding genes implicated in ribosome biogenesis (Qu et al., 2015). The *lncCOBRA1* promoter contained both a *Telo*-box and two Site II elements upstream of a TATA-box in the *lncCOBRA1* promoter, suggesting it is regulated in a similar manner to canonical snoRNAs and may be coordinated with genes related to ribosome biogenesis (**Supplemental Figure 3C**). In addition, the promoter contained a conserved non-coding sequence (CNS) (Velde et al., 2014), which are shown to be highly associated with genes encoding transcription factors and developmental genes and are enriched for transcription factor binding sites (Burgess and Freeling, 2014). The presence of a CNS further emphasizes the conservation of the *lncCOBRA1* gene locus (**Supplemental Figure 3C**). Overall, *lncCOBRA1* is a highly conserved lincRNA that is trimmed at its 3’ end post-transcriptionally to generate a ∼500-600 nt lincRNA.

### Loss of *lncCOBRA1* results in delayed germination and smaller plants

To examine the function of *lncCOBRA1*, we obtained a T-DNA insertion line (*lnccobra1-1;* SALK_086689) from the Arabidopsis Biological Resource Center with an insertion upstream of sno-*COBRA1A* and generated a complete *lncCOBRA1* null (*lnccobra1-2*) using CRISPR gene editing (**Figure 3A**). PCR and Sanger sequencing confirmed that the CRISPR guide RNAs caused a large deletion of 1325 bp (**Supplemental Figure 4A-B**). This larger than expected deletion was likely a product of double strand break repair (Korablev et al., 2020) and importantly did not disrupt the surrounding genes.

**Figure 3:**
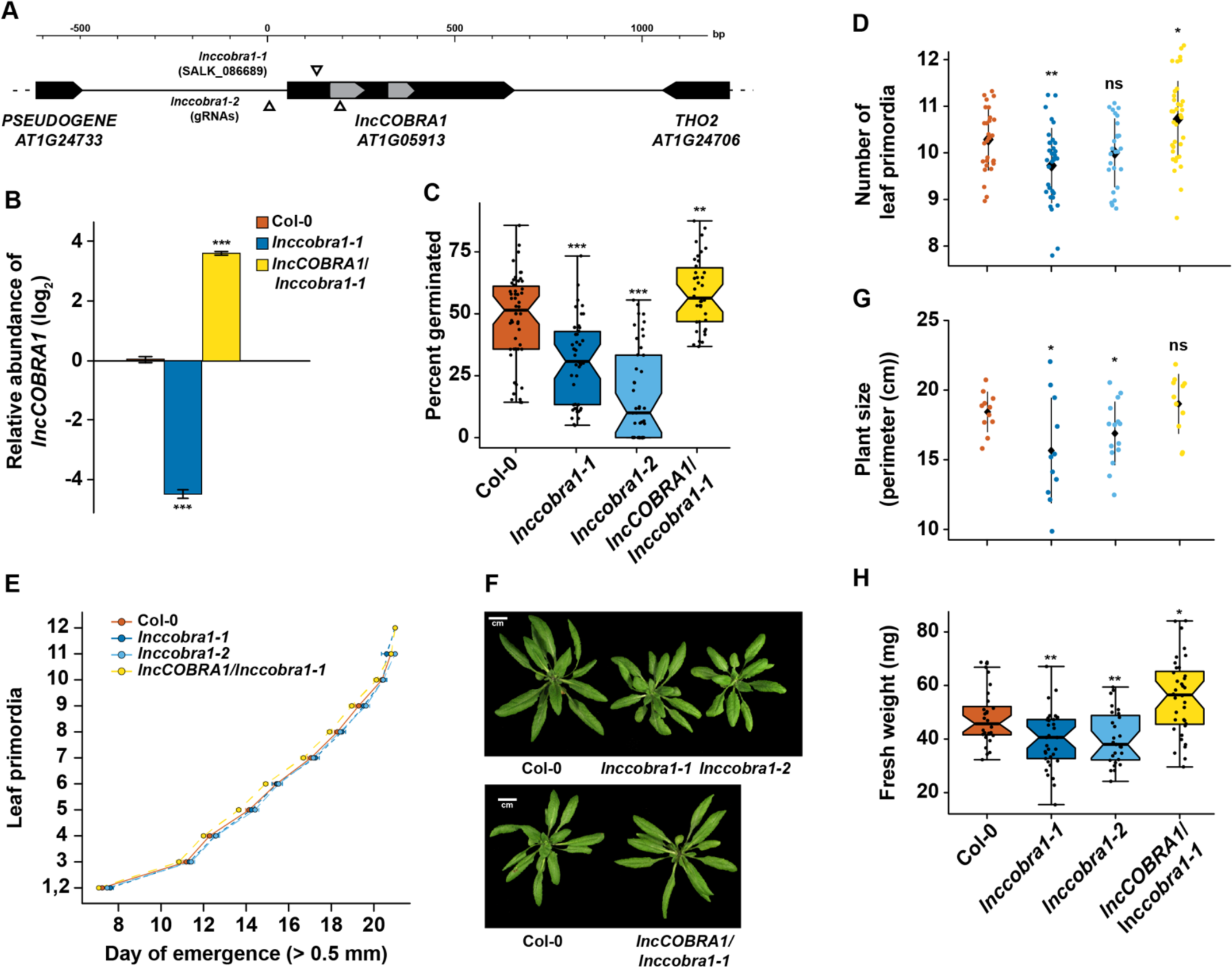
Loss of *lncCOBRA1* results in delayed germination and smaller plants. (A) Diagram of *lncCOBRA1* (*AT1G05913*) locus. Gray arrows represent the two snoRNAs annotated within *lncCOBRA1*. Triangles represent the location of the T-DNA insertion in SALK_086689 and location of the two guide RNAs used to generate a CRISPR deletion. (B) Relative abundance of *lncCOBRA1* in Col-0, *lnccobra1-1,* and *COBRA1*/*lnccobra1-1*. Abundance is normalized by the geomean of *UBC9* and *UBC10* and relative to Col-0. *** denotes p-value < 0.001; Wilcoxon t-test. N = 3. Error bars represent SEM. (C) Percent of seeds germinated 48 hr after sowing. Over 600 seedlings were measured per genotype on over 37 independent plates. *** denotes p-value < 0.001; Wilcoxon t-test. (D) Number of leaf primordia > 0.5 mm in 3-week-old plants. N > 27 plants per genotype. ns,*, and ** denotes p-value > 0.05, < 0.05, and < 0.01, respectively; Wilcoxon t-test. Black diamond represents the mean +/- SD. (E) Leaf initiation rate. The date was recorded for the first day each leaf primordia was visible by eye, ∼0.5 mm. N > 27 plants per genotype. Error bars represent SEM. (F) Representative images of 5-week-old Col-0, *lnccobra1-1*, *lnccobra1-2*, and *lncCOBRA1/lnccobra1-1*. Plants were grown in a 16/8 hr light/dark photoperiod at 22°C. All photos were taken the same day. (G) Plant perimeter analysis using ImageJ (see Methods). N > 11 per genotype. * denotes p-value < 0.05; Wilcoxon t-test. Black diamond represents the mean +/- SD. (H) Fresh weight of aerial tissue from 3-week-old plants. N > 27 plants per genotype. * and** denote p-value < 0.05 and < 0.01, respectively; Wilcoxon t-test.

*lnccobra1-1* had significantly (p-value < 0.001; Wilcoxon t-test) depleted levels of *lncCOBRA1* as measured by qRT-PCR and *lnccobra1-2* levels were unmeasurable as it is a null mutant with the entire gene deleted (**Figure 3B** and **Supplemental Figure 4C**) Levels and processing of rRNAs were minimally affected (**Supplemental Figure 4D-E**). Furthermore, the T-DNA insertion and CRISPR deletion were specific for decreasing *lncCOBRA1* as levels of the downstream protein-coding gene, *THO2,* were mostly unaffected in either mutant line (**Supplemental Figure 4D**). We did identify a slight but significant increase in *5.8S rRNA*, *18S rRNA*, and *25S rRNA* levels, but no visible changes in rRNA processing in the mutants compared to Col-0 (**Supplemental Figure 4E**). Thus, *lncCOBRA1* likely does not influence rRNA processing even though it contains two well-conserved snoRNA domains.

We also complemented the *lnccobra1-1* mutant by introducing the entire genomic region between the two neighboring genes into this genetic background (*lncCOBRA1pro::lncCOBRA1* / *lnccobra1-1*; *lncCOBRA1/lnccobra1-1*). *lncCOBRA1* complementation resulted in a significant increase in *lncCOBRA1* levels (**Figure 3B** and **Supplemental Figure 4C**). This overexpression of *lncCOBRA1* eliminated the slight but significant increase in *5.8S rRNA*, *18SrRNA*, *and 25S rRNA* levels observed in the mutant alleles (p-value > 0.05; Wilcoxon t-test), suggesting that the slight increase in abundance may in fact be due to the loss of *lncCOBRA1* (**Supplemental Figure 4D**). In total, our findings indicate that both *lnccobra1* mutant lines specifically and significantly decrease the levels of this lincRNA.

Given the high abundance of *lncCOBRA1* during seed germination (**Figure 1**), we examined cotyledon emergence of Col-0, *lnccobra1-1*, *lnccobra1-2*, and *lncCOBRA1/lnccobra1-1* seeds 48 hours after sowing as a proxy for germination defects. We observed that significantly (p-value < 0.001; Wilcoxon t-test) fewer *lnccobra1-1* and *lnccobra1-2* seeds germinated than in the Col-0 background, while significantly (p-value < 0.01; Wilcoxon t-test) more *lncCOBRA1/lnccobra1-1* seeds germinated at 48 hours (**Figure 3C**), suggesting that *lncCOBRA1* levels affect seed germination.

The effects of *lncCOBRA1* on germination persisted throughout vegetative growth, as 3-week-old *lnccobra1-1* plants were slightly but significantly (∼0.5 leaves; p-value < 0.01; Wilcoxon t-test) delayed in leaf production compared to same aged Col-0 plants. This same trend was also observed in *lnccobra1-2* plants, but not to a level of statistical significance (p-value > 0.05; Wilcoxon t-test) (**Figure 3D**). Increased levels of *lncCOBRA1* in *lncCOBRA1/lnccobra1-1* plants led to significantly (∼0.5 leaves, p-value < 0.05; Wilcoxon t-test) more leaves than Col-0 (**Figure 3D**), suggesting *lncCOBRA1* is responsible for this phenotype. This change in number of leaves at 3-weeks after planting was not due to a change in the overall growth rate of the plants, as there is no change in rate of leaf initiation in *lnccobra1-1*, *lnccobra1-2,* or *lncCOBRA1/lnccobra1-1* compared to Col-0 (**Figure 3E**). *lnccobra1-1* and *lnccobra1-2* plants were also substantially smaller than Col-0 plants, while the plants overexpressing *lncCOBRA1* (*lncCOBRA1/lnccobra1-1*) rescued this phenotype and resulted in plants that were slightly larger in both 3- and 5-week-old plants (**Figures 3F-H** **and Supplemental Figure 5A**). Aside from overall size of the plants, the individual rosette leaves were also smaller in the mutant plant lines (**Supplemental Figure 5B**). Since altered *lncCOBRA1* levels did not affect the rate of growth (**Figure 3E**), it is possible that the smaller nature of *lnccobra1-1* and *lnccobra1-2* may be due to a change in either the number or size of leaf cells, though this needs to be probed further. Overall, levels of *lncCOBRA1* effect seed germination, and these germination effects persist through vegetative growth, resulting in plants that are smaller or larger than Col-0 when *lncCOBRA1* levels are decreased or increased, respectively.

### *lncCOBRA1* interacts with a wide variety of proteins

To begin to understand the molecular function of *lncCOBRA1,* we set out to identify what proteins bind *lncCOBRA1*, as *lncCOBRA1* was initially identified for having sites of RBP binding (**Supplemental Figure 1A**) (Gosai et al., 2015). To do so, we performed chromatin isolation by RNA purification followed by mass spectrometry (ChIRP-MS) (Chu et al., 2015). In this technique, we incubated lysates from 5-day-old Col-0 and *lnccobra1-2* seedlings with biotinylated probes antisense to *lncCOBRA1* (**Figure 4A**) or a scrambled sequence as a negative control. We then used streptavidin coated beads to pull down *lncCOBRA1*, isolated proteins bound, and performed mass spectrometry. We confirmed the efficacy of the pulldown by qRT-PCR and found *lncCOBRA1* was significantly (p-value < 0.001; Wilcoxon t-test) enriched with probes antisense to *lncCOBRA1* compared to the scrambled sequence control probes, indicating that the *lncCOBRA1* probes are highly specific (**Figure 4A**). Importantly, enrichment of *lncCOBRA1* with the experimental probes was significantly (p-value < 0.001; Wilcoxon t-test) depleted when ChIRP was performed in *lnccobra1-2* null seedlings, (**Figure 4A**). As *lncCOBRA1* contains two snoRNA domains, we also asked whether *lncCOBRA1* directly interacted with rRNAs and found that *lncCOBRA1* probes did not enrich for *5.8S rRNA*, *18S rRNA*, or *25S rRNA* relative to scrambled sequence control probes (**Supplemental Figure 6A**). This indicated that *lncCOBRA1* does not interact with rRNA, further confirming that the snoRNA domains within *lncCOBRA1* do not function like canonical snoRNAs (**Supplemental Figure 6A**).

**Figure 4:**
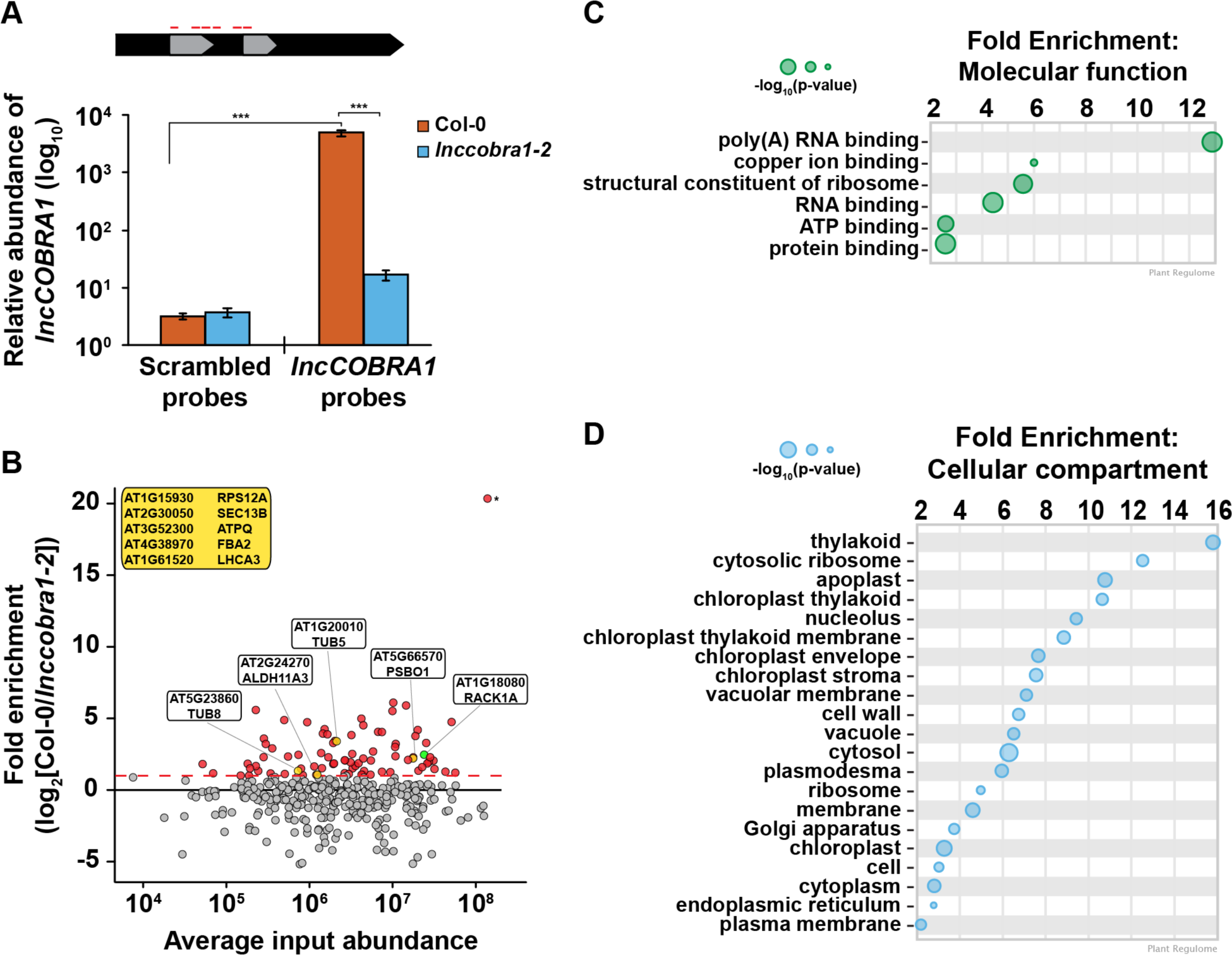
ChIRP enriches for *lncCOBRA1* and identifies 113 *lncCOBRA1*-interacting proteins. (A) Relative abundance of *lncCOBRA1* in ChIRP-MS experiments. Abundance normalized by *U6* and is relative to Col-0 input. Error bars represent SEM. ns,*,**, and *** denotes p-value > 0.05, < 0.05, < 0.01, and < 0.001, respectively; Wilcoxon t-test. N = 3. (B) Proteins identified from ChIRP followed by MS. X-axis is the average protein abundance in Col-0 and *cobra1-2* inputs. Y-axis is fold enrichment in Col-0 relative to *lnccobra1-2* with *lncCOBRA1* probes. All dots were enriched with *lncCOBRA1* probes compared to scrambled sequence probes. Red dots indicate proteins enriched with *lncCOBRA1* probes over scrambled and enriched at least 1-log2 fold change in Col-0 compared to *lnccobra1-2*. Green dot represents RACK1A. Yellow dots represent *lncCOBRA1*-interacting proteins that were experimentally shown to interact with RACK1A. Yellow box contains *lncCOBRA1*-interacting proteins that were experimentally shown to interact with RACK1A but were not identified in Col-0 with scrambled probes. N = 3. (C-D) Gene ontology enrichment analysis for molecular function (C) and cellular compartment (D) using Plant Regulomics (Ran et al., 2020) for *lncCOBRA1*-interacting proteins. Size of circles represents -log_10_(p-value).

After mass spectrometry, we set out to identify high confidence interactors from the ∼2200 proteins identified (**Supplemental Data Set 1A**). To do so, we required that proteins must be (1) identified in at least 2 out of 3 biological replicates of the *lncCOBRA1* pulldown in Col-0 plants (N = 469 proteins) (**Supplemental Data Set 1B**), (2) enriched with the *lncCOBRA1* probes compared to scrambled sequence control probes (N = 206), and (3) enriched > 2-fold in Col-0 compared to *lnccobra1-2* seedlings (N = 74; **Figure 4B**, red dots, and **Supplemental Figure 6B**). A total of 74 proteins were identified from these filtering steps. An additional 39 proteins were identified in at least 2 biological replicates in the *lncCOBRA1* pulldown but absent from control pulldowns (scrambled or *lnccobra1-2* background; **Table 2**). In total, 113 proteins were identified as high-confidence *lncCOBRA1*-interacting proteins, and specifically bound to *lncCOBRA1* in 5-day-old Col-0 seedlings.

*lncCOBRA1*-interacting proteins were significantly enriched for proteins with molecular function of RNA binding, and 37.5% (p-value < 5.21 x 10^-40^; hypergeometric test) were demonstrated to bind to RNA in a recent study identifying the RNA binding proteome of Arabidopsis leaves (Bach-Pages et al., 2020), supporting the claim that these proteins interact directly with *lncCOBRA1* (**Figure 4C** and **Supplemental Figure 6C**). Those proteins not demonstrated to have RNA binding capabilities may still interact with *lncCOBRA1* indirectly. In addition, several proteins involved in transcription regulation were identified, including PUR ALPHA-1 (PUR*α*), which has hypothesized roles in rRNA transcription (**Table 3**; Trémousaygue et al., 2003).

*lncCOBRA1*-interacting proteins were involved in a wide-range of biological functions, including response to cytokinin and abscisic acid (ABA), gluconeogenesis, and photorespiration (**Supplemental Figure 6D**) (Ran et al., 2020). Additionally, *lncCOBRA1-*interacting proteins were enriched for proteins functioning in ‘structural constituents of the ribosome’ and located in the cytoplasmic ribosome, chloroplasts, and the nucleolus (**Figure 4D**). In fact, twelve of the *lncCOBRA1-*interacting proteins (10.6%; p-value < 2.7 x 10^-13^; hypergeometric test; **Table 4**) were identified in a previous study identifying the nucleolar proteome (Pendle et al., 2005). The nucleolus is a non-membrane bound nuclear structure that is the site for ribosome assembly and maturation. Given the snoRNA domains in *lncCOBRA1* and the identification of cytoplasmic ribosomal constituents bound to the nuclear localized *lncCOBRA1*, we hypothesize that *lncCOBRA1* may be localized to the nucleolus, though this needs to be directly tested.

Aside from the nucleolus, *lncCOBRA1*-interacting proteins were also enriched in various chloroplast related compartments (**Figure 4D**). In fact, many *lncCOBRA1*-interacting proteins were localized in the chloroplast, including PSAD-II, PTAC16, TIG and RNASE J (RNJ; **Supplemental Data Set 1B**). Among these RNA binding *lncCOBRA1*-interacting proteins is RNaseJ (RNJ), which is the most enriched protein bound to *lncCOBRA1* in Col-0 relative to *lnccobra1-2* (**Figure 4B**; marked by asterisk). RNJ is a metallo-beta-lactamase protein that possesses endo- and 5’-3’ exonuclease activities in bacteria and chloroplasts within plants and is required for embryo and chloroplast development (Halpert et al., 2019) with roles in rRNA maturation and 5’ stability of mRNAs in bacteria (Mathy et al., 2007).

### *lncCOBRA1*-interacting proteins are highly interconnected

As proteins tend to act in complexes and *lncCOBRA1*-interacting proteins were enriched for proteins involved in protein binding (**Figure 4C**), we next asked if there were known interactions among the 113 high-confidence *lncCOBRA1*-interacting proteins (**Figure 4B; Table 2**). Using STRING, we generated a protein-protein interaction (PPI) network which formed significantly (p-value < 1.0 x 10^-16^; STRING) more interactions than expected, indicating that *lncCOBRA1*-interacting proteins had more interactions among themselves than what would be expected for a random set of proteins of a similar size from the Arabidopsis proteome **(Supplemental Figure 7A**) (Szklarczyk et al., 2019). Using k-means clustering, the proteins within the network were further grouped into 5 clusters (green, cyan, blue, red, and yellow) (**Supplemental Figures 7A-B**). Each cluster represented distinct groups of proteins with cytokinin response-related and photosynthetic proteins, glycolytic proteins, and mRNA splicing-related proteins clustering together to form the green, cyan and blue clusters, respectively (Huang et al., 2009). Of the five clusters, blue, green, and cyan were interlaced throughout the network, and hard to distinguish between each other. The red cluster was the most spread out, lying on the periphery of the network with very little significant enrichment for biological processes or cellular compartments, indicating this cluster represents a variety of different proteins with a range of functions (**Supplemental Figure 7B)**.

Within the red cluster lies the trihelix DNA binding transcription factor 6B-INTERACTING PROTEIN 1-LIKE (ASIL1) (**Supplemental Figure 7A**), which was previously shown to be involved in repressing seed maturation genes during seed germination and seedling development (Gao et al., 2009) and was also previously identified in the nucleolus (**Table 2**).

Since numerous nuclear lincRNAs function in gene regulation by binding and directing transcription factors to the correct genomic loci, and ASIL1 regulates germination, which is mis-regulated in *lnccobra1-1* and *lnccobra1-2* plants, it is possible that *lncCOBRA1* interacts with ASIL1 to affect seed maturation genes during seed germination and seedling development, but further studies are required to test this.

A closer examination of the yellow cluster, which was the most compact group (**Figure 5A**), revealed that this close network was enriched for proteins involved in ribosome biogenesis, rRNA processing, response to cytokinin, RNA binding, and constituents of the ribosome (**Figures 5B-D; Supplemental Figure 7B**). This cluster was also enriched for proteins localized in the nucleolus and ribosome (**Figures 5B-D** and **Supplemental Figure 7B**). A major node within the yellow cluster was RECEPTOR FOR ACTIVATED C KINASE 1A (RACK1A; encoded by *ATARCA*) (**Figure 5A**). RACK1A is a major subunit of RACK1, which is a highly conserved scaffold protein present in all eukaryotic organisms studied, from *Chlamydomonas* to plants and humans (Adams et al., 2011). Several proteomics studies have identified a total of 293 proteins that interact with RACK1A (Stark et al., 2006; Cheng et al., 2015; Kundu et al., 2013; Guo et al., 2019; Olejnik et al., 2011; Speth et al., 2013), 40 of which (13.7%; p-value < 2.1 x 10^-28^; hypergeometric test) were identified in at least 2 biological replicates of *lncCOBRA1* pulldown in Col-0 (**Figure 5E**). This included RACK1B, another major subunit of RACK1 (Guo and Chen, 2008). Nearly 25% of the identified RACK1A-interacting proteins that were identified in ChIRP were specifically bound to *lncCOBRA1* in Col-0 compared to *lnccobra1-2* (N = 9; **Figure 5B**, yellow dots), providing evidence that *lncCOBRA1* interacts with RACK1A.

**Figure 5:**
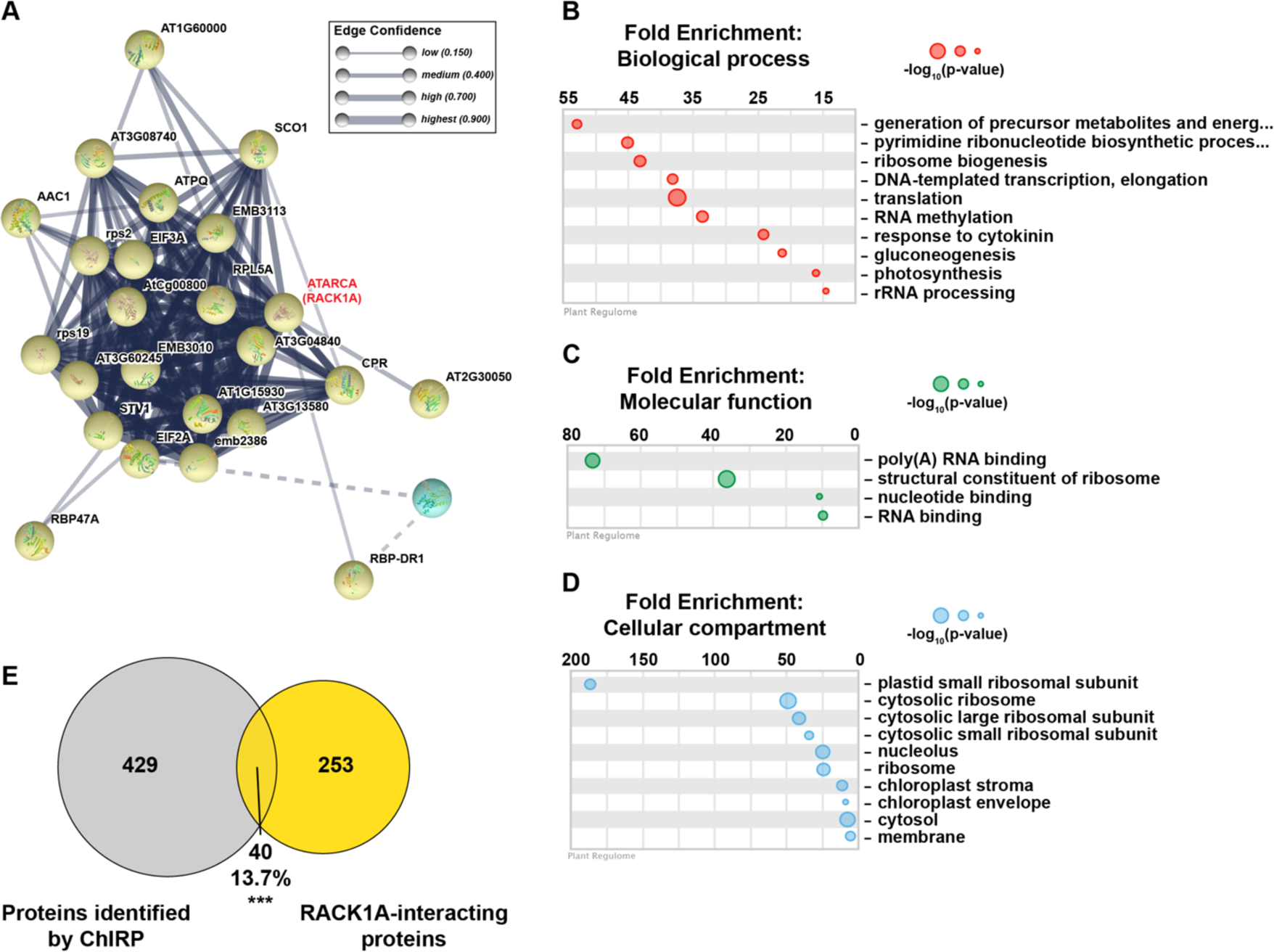
*lncCOBRA1* interacts with RACK1A and a tight network of proteins related to ribosome biogenesis. (A) Yellow protein-protein interaction k-means cluster generated from STRING (Szklarczyk et al., 2019). Thickness of lines connecting notes indicates the confidence of that protein-protein interaction. Dotted line indicates interaction with a different cluster (See Supplemental Figure 7 for full network). (B-D) Gene ontology enrichment analysis for biological process (B), molecular function (C), and cellular compartment (D) using Plant Regulomics (Ran et al., 2020) for *lncCOBRA1*-interacting proteins in the yellow cluster. Size of circles represents -log_10_(p-value). (E) Overlap between proteins identified in at least two biological replicates of Col-0 ChIRP with the *lncCOBRA1* probes and proteins identified as RACK1A binding. *** denotes p-value < 0.001; Hypergeometric test.

## DISCUSSION

In this study, we use genetic, biochemical, and proteomic analyses to describe a highly conserved, previously uncharacterized sno-lincRNA with functions in seed germination and development. We reveal that *lncCOBRA1* is a ∼500-600 nt lincRNA with germination-, developmental-, and tissue-specific patterns of abundance, with high abundance early during seed germination and decreases as development progresses. Further, we demonstrate that loss of *lncCOBRA1* results in delayed cotyledon emergence and overall smaller plants. We demonstrate that *lncCOBRA1* interacts with a wide variety of proteins, including many nucleolar proteins and scaffold proteins, including the highly conserved RACK1 subunit RACK1A. Though additional experiments are needed, we hypothesize that *lncCOBRA1* may act as a scaffold to bring together proteins involved in several different processes to ultimately regulate plant germination and development.

### Identification of highly conserved, protein-bound nuclear lincRNAs from transcriptome-wide analyses

Here, we describe a set of lincRNAs named *CONSERVED IN* BRASSICA RAPA *1-14* (*lncCOBRA1-14*) that were identified for their interactions with nuclear RBPs and sequence conservation in *Brassica rapa* (**Supplemental Figure 1A**) (Gosai et al., 2015). Of the 14 *lncCOBRA* transcripts profiled, 9 contained one or more snoRNAs annotated within it, revealing a previously unidentified class of lincRNAs containing snoRNAs (sno-lincRNAs) in Arabidopsis (**Table 1**). SnoRNAs are a family of conserved nuclear small RNAs (70 - 200 nt) that are usually concentrated in the Cajal bodies or nucleolus. They traditionally function to modify rRNA or participate in the processing and maturation of ribosomal subunits, where binding of core nucleolar proteins protects the mature snoRNAs and aids in proper function (Rodor et al., 2010).

We predict that the presence of snoRNA sequences in these lincRNAs likely results in their interaction with RBPs, as the annotated snoRNA domains overlap with the protein-bound sites identified previously, and snoRNA sequences are known to be highly protein-bound. Additionally, since snoRNAs are nuclear retained (**Figure 1**), we predict that the snoRNA sequences contained in these *lncCOBRA* transcripts permit their nuclear retention, though future experiments are needed to test this hypothesis. Most *lncCOBRA* transcripts demonstrated specific patterns of abundance during seed germination. Interestingly, *COBRA* lincRNAs that lacked snoRNA sequences demonstrated the least specificity in abundance patterns during germination (*lncCOBRA8, 9*, *13*, and *14*) (**Supplemental Figure 1B**). Ultimately, this suggests that sno-lincRNAs may be important for germination in Arabidopsis, while conserved, protein-bound lincRNAs that lack snoRNAs may function in different biological processes.

In mammals, the majority of functional snoRNAs are encoded within introns and processed from excised and debranched introns by exonucleolytic trimming. Similarly, all identified mammalian sno-lncRNAs are generated from excised introns as well (Xing and Chen, 2018). In Arabidopsis, while identified snoRNAs in Arabidopsis appear to be homologs of yeast and animal counterparts, they are not encoded within introns but are instead primarily transcribed from intergenic regions as polycistronic gene clusters. As such, the *lncCOBRA* sno-lincRNAs described here are also transcribed from intergenic regions throughout the genome. Thus, *lncCOBRA* sno-lincRNAs represent a previously uncharacterized class of lincRNAs with potentially important biological functions that warrant future studies.

### Regulation of lncCOBRA1 transcription

*lncCOBRA1* contains several conserved elements within its promoter known to be present in the promoters of genes involved in ribosome biogenesis and translation. This includes Telo-box and Site II elements (*Telo*SII) (**Supplemental Figure 3D**). Interestingly, the Telo-box is known to be bound by the *lncCOBRA1*-interacting transcription factor PUR ALPHA-1 (PUR*α*) (**Table 2-3**) (Tremousaygue et al., 1999). PUR*α* is a homolog of the animal nuclear protein PUR ALPHA (*PURA*) which is a member of the sequence-specific single-stranded nucleic acid-binding Pur family of proteins. The amino acid sequence of Pura is extraordinarily conserved in sequence from bacteria through humans, where it functions as a transcriptional activator, and as an RNA transport protein. While less is known about PUR*α* in Arabidopsis, it was identified to be an RBP (Bach-Pages et al., 2020) and was previously demonstrated to interact with TEOSINTE BRANCHED 1, CYCLOIDEA, PCF (TCP)-DOMAIN FAMILY PROTEIN 20 (TCP20) (Trémousaygue et al., 2003). TCP20 also binds *Telo*SII elements and regulates expression of ribosomal protein genes (Trémousaygue et al., 2003). In Arabidopsis, nearly all ribosomal protein genes and other genes involved in ribosome biogenesis and translation contain *Telo*SII elements in their promoters (Gaspin et al., 2010). This combined *Telo*SII element is found upstream of the TATA box and acts to coordinate expression of snoRNAs and ribosome biogenesis (Qu et al., 2015). Thus, the interaction between PUR*α* and *lncCOBRA1* could suggest the *lncCOBRA1* binds to PUR*α* to regulate its own expression. Additionally, the presence of the *Telo*SII elements in the *lncCOBRA1* promoter suggests that *lncCOBRA1* may be expressed in a coordinated manner with ribosomal proteins, implicating it in ribosome biogenesis.

### lncCOBRA1-interacting proteins may mediate germination phenotype observed in mutants

RACK1 is a versatile scaffold protein that can bind to numerous signaling molecules from diverse signal transduction pathways (Guo et al., 2007). In Arabidopsis, RACK1 plays an important role in maintaining 60S ribosome biogenesis and 80S monosome assembly, as *rack1a rack1b* double mutants have a decrease in abundance of the 60S ribosomal subunit and 80S monosomes, but no differences in polysomes, suggesting a role for RACK1 in ribosome biogenesis (Guo et al., 2011). Since RACK1A interacts with ribosomal proteins, generally affects translation and responds to several hormones, this suggests that RACK1 has a dual role in signaling and translation, as observed previously for the RACK1 homolog in mammals (Guo et al., 2011).

Additionally, mutants in RACK1A had smaller rosette leaf size and delayed flowering and leaf development under short day conditions (8/16 hr photoperiod) (Chen et al., 2006). When grown under long day conditions (16/8 hr photoperiod), many of the strong phenotypes observed under short day were alleviated and *rack1a* plants grew at similar rates to wild type, but had slightly smaller rosette leaf size, a phenotype that was exacerbated when additional subunits of RACK1 were deleted (Wang et al., 2019). Overall, *rack1a* plants grown under long day conditions appear to phenocopy *lnccobra1* mutants, potentially suggesting a functional link between RACK1A and *lncCOBRA1*. Moreover, *rack1a* mutants were hypersensitive to ABA, suggesting a role of RACK1A in negatively regulating ABA-mediated seed germination and development (Chen et al., 2006; Guo et al., 2009, 2011). Ultimately, the hypersensitivity of *rack1a* to ABA suggests that RACK1A negatively regulates ABA-mediated seed germination and development.

Given the evidence of RACK1-*lncCOBRA1* interaction (**Figures 4-5**) along with similarities in the phenotype of null mutants (Chen et al., 2006; Guo et al., 2019) and protein binding partners (**Figures 4-5**), we hypothesize that *lncCOBRA1* functions with RACK1A as a scaffold to regulate plant germination and development.

### RNASE J is the highest enriched protein-bound to lncCOBRA1

The protein with the highest enrichment for *lncCOBRA1* binding in Col-0 relative to *lnccobra1-2* was RIBONUCLEASE J (RNASE J; RNJ) (**Figure 4B**). *RNJ* encodes a metallo-beta-lactamase protein that possesses endo- and 5’-3’ exonuclease activities in bacteria and chloroplasts within plants and is required for embryo and chloroplast development (Halpert et al., 2019). While RNase J plays important roles in rRNA maturation and 5’ stability of mRNAs in bacteria (Mathy et al., 2007), it does not function in the cleavage of polycistronic rRNAs or mRNA precursors in Arabidopsis (Sharwood et al., 2011). Instead, loss of RNase J resulted in a massive accumulation of antisense RNAs, suggesting that RNase J is responsible for degradation of these RNAs generated by the inability of chloroplast RNA polymerase to terminate transcript effectively. The antisense RNAs would otherwise form duplexes with sense strand transcripts and prevent translation (Sharwood et al., 2011). While RNase J is described to be chloroplast localized, it is also predicted to be located in the nucleus by computational predictions (Kaundal et al., 2010). Further, previously, we previously identified a protein thought to be solely chloroplast localized in the nucleus (Gosai et al., 2015). Thus, it is possible that RNase J is in the nucleus, though this needs to be directly experimentally validated.

RNases are essential for non-coding RNA processing and each RNase can have a multitude of targets. For example, RNase P is an endoribonuclease canonically functions to process the 5’ termini of pre-tRNAs but can also cleave other tRNA like structures in the 3’ end of lncRNAs to form mature 3’ ends (Wilusz et al., 2008, 2011; Sunwoo et al., 2009). Additionally, RNase mitochondrial RNA processing (MRP) was originally identified as an RNA-protein endoribonuclease that processes RNA primers of DNA replication in the mitochondria but is actually predominantly found in the nucleolus where it participates in pre-rRNA processing (Lee et al., 1996). Thus, it is possible that RNase J possesses additional functions than previously described, possibly mediated by its interaction with *lncCOBRA1*. Given its function in ribosome maturation in bacteria and the multiple functions of RNases on ncRNAs described previously, we posit that RNase J may have additional function in sno-lincRNA processing in Arabidopsis, specifically the 3’ end processing we observe for *lncCOBRA1*, but future studies will be required to support this hypothesis.

In total, using transcriptome-wide analyses we identified functional candidate lncRNAs based on sequence conservation and the presence of RBP binding sites. We further show the loss of *lncCOBRA1* results in growth phenotypes. While future studies are required, we provide evidence that *lncCOBRA1* interacts with a plethora of proteins involved in many different processes. Overall, we hypothesize that *lncCOBRA1* acts as a scaffold to bring together many different proteins to regulate normal biological processes, including ribosome biogenesis.

## AUTHOR CONTRIBUTIONS

M.C.K and B.D.G conceived the study. M.C.K, H.J.K, K.R.P, and B.D.G designed the experiments. M.C.K, H.J.K., and K.R.P. performed experiments and analyzed the data. M.C.K. and B.D.G. wrote the paper with assistance from all authors. The authors have read and approved the manuscript for publication.

## Supporting information

Supplemental Data Set 1

Supplemental Data Set 2

## ACKNOWLEDGMENTS

The authors would like to thank the members of the B.D.G. lab both past and present for helpful discussions. This work was funded by a Graduate Women in Science Fellowship National Fellowship to M.C.K, NSF grants IOS-1849708 and MCB-1623887 to B.D.G., IOS-1758532 to A.D.L.N., and IOS-1444490 to B.D.G, M.A.B., and E.L. The funders had no role in study design, data collection and analysis, decision to publish, or preparation of the manuscript.

## MATERIALS AND METHODS

### Plant materials and growth conditions

All plants were of the Columbia-0 ecotype and were grown in controlled chambers with a cycle of 16 hours light and 8 hours dark at 22°C. All seeds used for plate growth were sterilized in 100% ethanol for one-minute followed by a 10-minute wash with 30% Clorox and 0.01% Tween-20 solution and rinsed 5 times with sterilized water. Seeds were then plated and grown on ½ MS agar plates with 1% sucrose and 0.8% Phytoblend and stratified by cold treating at 4°C for 48 hrs then placed in growth chambers with the parameters noted above.

*lncCOBRA1* was previously referred to as *AT1NC031460* in Liu *et al*. and *AT1G05913* in the Araport11 genome annotation. *lnccobra1-1* (SALK_086689) was purchased from the Arabidopsis Biological Resource Center and backcrossed once to Col-0, segregated, and homozygous mutants obtained and validated by PCR. qRT-PCR was used to validate significant depletion in the abundance of *lncCOBRA1*.

### CRISPR/Cas9 plasmid construction and mutation identification

To generate *lnccobra1-2*, the suite of plasmids designed for multiplexed CRISPR genome editing by Lowder *et al*. (2015) were acquired from Addgene (https://www.addgene.org) and used to generate Arabidopsis CRISPR-Cas9 transformation vectors (Lowder et al., 2015). Two different guide RNAs were designed using the CRISPRdirect website (https://crispr.dbcls.jp) targeting *AT1G05913*. Because Cas9 was chosen to perform genome editing, 5’ -NGG-3’ was used as the protospacer adjacent motif (PAM) sequence requirement. The *Arabidopsis thaliana* TAIR10 genome was used to ensure the specificity of chosen guide RNAs. The first guide RNA (protospacer sequence: 5’ -TATGATTTGATCATCATCGG-3’) is located approximately 50 base pairs upstream of the *AT1G05913* transcription start site, and the second guide RNA (protospacer sequence: 5’ -TATATGGCTCTGGAAGAGGG-3’) is located approximately 121 base pairs downstream of the *AT1G05913* transcription start site. Complimentary oligos were designed for each protospacer that contained overhangs compatible with the Arabidopsis *U6* promoter driven guide RNA vectors designed by Lowder *et al*. (2015) (vectors pYPQ131-pYPQ134) (Lowder et al., 2015).

To generate a CRISPR-Cas9 transformation vector containing two guide RNAs targeting *AT1G05913*, the cloning procedures provided by Lowder *et al*. (2015) were followed (Lowder et al., 2015). Briefly, each protospacer sequence described above was annealed using complimentary oligos to create a double stranded DNA fragment and then ligated into the vectors pYPQ131 and pYPQ132, respectively. pYPQ131 and pYPQ132 with correctly inserted protospacer sequences were used in a Golden Gate assembly reaction with pYPQ142 to generate a Gateway-compatible entry vector. The pYPQ142 vector with both guide RNAs correctly inserted, along with pYPQ154 carrying an Arabidopsis codon optimized Cas9, and pUBQ10:GW (Stock CD3-1947 from the Arabidopsis Biological Resource Center) were used in a Gateway LR reaction (Thermo Fisher Scientific; Carlsblad, CA, USA) to generate the final transformation vector. The final vector was transformed into wild type *Arabidopsis thaliana* (Col-0) using the floral dip method (Clough and Bent, 1998).

Successful transformants were selected using Glufosinate-ammonium and allowed to set seed to acquire second generation transformants (T2). T2 plants were genotyped to test for a deletion in *AT1G05913* using the PCR primers 5’ – CGCTTGTTCAACTCCAAAAAG-3’ and 5’-TTTTGGTATATAAGCTGATGGC-3’. A large band shift was detected in one T2 plant (wild type product size: 1,600 bp, observed product size: approximately 200 bp) (**Supplemental Figure 4A**), and Sanger sequencing confirmed the deletion to be 1,325 bp. All primers are listed in **Supplemental Data Set 2**.

### Plasmid construction and generation of *lncCOBRA1/lnccobra1-1*

To generate *lncCOBRA1* promoter:: *lncCOBRA1/lnccobra1-1*, the entire 1509 bp between the two neighboring genes was amplified from Col-0 genomic DNA and cloned into BspEI and BstEII restriction enzyme sites of pCAMBIA3301. Transgenic plants were obtained and selected as previously described (Zhang et al., 2006). All primers are listed in **Supplemental Data Set 2**.

### RNA extraction

RNA was extracted from the tissues denoted using a liquid nitrogen cooled mortar and pestle. Ground, frozen tissue was transferred to Qiazol lysis reagent (Qiagen; Valencia, CA, USA) and further homogenized using QIAshredders (Qiagen; Valencia, CA, USA). RNA was then isolated using the miRNeasy mini columns as described by the manufacturers’ protocol (Qiagen; Valencia, CA, USA). Following elution from the miRNeasy column, RNA was treated with RNase-free DNase (Qiagen, Valencia, CA, USA) for 25 minutes at room temperature, ethanol precipitated and resuspended in nuclease-free water supplemented with 1.25% RNaseOUT (Life Technologies; Carlsbad, CA, USA).

### qRT-PCR

All reverse transcription (RT) reactions were performed using SuperScript II following the manufacturers’ instructions with 2.5 mM Random Hexamers (Qiagen; Valencia, CA, USA), 100 units SuperScript II and 30 units RNaseOUT (Invitrogen; Carlsbad, CA, USA) for 2 minutes at 25°C, 90 minutes 42°C, 5 minutes 95°C, hold at 4°C. Before qRT-PCR, cDNA was diluted 1:10 for all qRT-PCR reactions except for ChIRP in which the RT reaction was diluted 1:5.

qRT-PCR was performed with 2X SYBR Green qRT-PCR Master Mix with Rox #2 (Bimake; Houston, TX, USA), as follows per well: 10 μL 2X SYBR Green Master Mix, 1.5 μL cDNA (diluted 1:10), 0.4 μL Rox #2. 2.1 μL water, 6 μL combined 1.5 μM forward and reverse primers. All qRT-PCR reactions were performed in 3 technical replicates and all primers tested using water to detect background signal and melt curves were analyzed for a single peak. All qRT-PCRs were run using the following program: 95°C for 10 minutes; 40 cycles of 95°C 30 sec, 55°C 30 sec, 72°C 30 sec. Melt curves were generated by heating the final PCR 1.6°C/s to 95°C for 15 sec, decreasing the temperature to 60°C at 1.6°C/s and slowly increasing back to 95°C at 0.1°C/s. Unless otherwise noted, all qRT-PCR experiments were normalized to the geomean of *UBC9* and *UBC10*. All primers are listed in **Supplemental Data Set 2**.

### INTACT

To examine RNA abundance in nuclei and cytoplasmic fractions, seeds ubiquitously expressing a biotin ligase receptor peptide fusion protein that is targeted to the nuclear envelope (UBQ10:NTF/ACT2p:BirA Columbia-0 ecotype) were used (Deal and Henikoff, 2010, 2011). After 7 days, seedlings were collected, and flash frozen in liquid nitrogen and stored at - 80°C for further processing. The isolation of nuclei tagged in specific cell types (INTACT) (Deal and Henikoff, 2010, 2011) technique was used to isolate pure nuclear and cytoplasmic fractions and RNA extracted before RT and qPCR as described above.

### Tissue collection

For the germination time course, seedlings were collected 2, 3, 4, 5, 7, and 10 days after stratification and flash frozen in liquid nitrogen and stored in -80°C for further processing.

Tissues from 5-week-old Col-0 plants were collected, flash frozen in liquid nitrogen, and stored in -80°C until processing for examining the tissue specificity of *lncCOBRA1* abundance. The sample of adult leaves included a mix of rosette leaves older than leaves 1-4 which were denoted juvenile leaves.

### Brassicaceae *lncCOBRA1* sequence alignments

To identify putative sequence homologs of the *AT1G05913* gene, the entire Arabidopsis cDNA sequence was used as query for BLAST using CoGeBlast (https://genomevolution.org/CoGe/CoGeBlast.pl) using default parameters (E-value: 1e-5, Word size: 8, Gap Costs: Existence-5 Extension-2, Match/Mismatch Scores: 1,-2) against representative Brassicaceae species. The top hits for each species were selected based on e-value and quality score and used for subsequent sequence alignments. Selected sequences were aligned using Geneious Prime (Geneious | Bioinformatics Solutions for the Analysis of Molecular Sequence Data, 2019) with the Multiple Alignment tool, utilizing the Geneious Alignment default parameters (Alignment type: Global alignment with free end gaps, Cost Matrix: 70% similarity, Gap open penalty: 12, Gap extension penalty: 3, Refinement iterations: 2).

### Rapid Amplification of cDNA Ends (RACE)

#### 5’ RACE

5 μg of RNA from 5-day-old seedlings was first treated with 1 unit of Shrimp Alkaline Phosphatase (SAP; USB Products, Affymetrix, Inc.; Cleveland, OH, USA) in 1X SAP buffer provided and supplemented with 1 mM DTT and 60 units RNaseOUT (Invitrogen; Carlsbad, CA, USA) for 1 hr at 37°C. The SAP reaction was inactivated for 15 minutes at 65°C and the RNA ethanol precipitated overnight. To remove any 5’ m^7^G caps, 500 ng of the SAP-treated RNA was treated with 12.5 units RNA 5’ Pyrophophohydrolase (RppH; New England BioLabs; Ipswitch, MA, USA) in 1X T4 RNA Ligase Buffer (New England BioLabs; Ipswitch, MA, USA) supplemented with 20 units RNaseOUT (Invitrogen; Carlsbad, CA, USA) in a total reaction volume of 10 μL for 1 hr at 37°C and stored at -20°C overnight.

On the following day, the 5’ adapter was added. To the 10 μL RppH reaction, we added 1 μL of 5’ RNA adapter (25 μM; RA5; 5’-GUUCAGAGUUCUACAGUCCGACGAUC -3’) that was first heated to 70°C for 2 minutes followed by 2 minutes on ice to relieve secondary structures, 1 μL 10 mM ATP (New England BioLabs; Ipswitch, MA, USA), 10 units T4 RNA Ligase 1 (New England BioLabs; Ipswitch, MA, USA), 1 μL T4 RNA Ligase Buffer (New England BioLabs; Ipswitch, MA, USA), and 40 units RNaseOUT (Invitrogen; Carlsbad, CA, USA) and incubated for 3 hrs at 20°C followed by an overnight ethanol precipitated. For cDNA synthesis, 1 μL of gene specific primer (10 μM; “*lncCOBRA1* qPCR Reverse set 2”) was added to the ligase reaction and heat treated at 80°C for 3 minutes followed by 2 minutes on ice. Reverse transcription was performed with 100 units SuperScript II in 1X First Strand Buffer, 2 mM dNTPs, 10 mM DTT, and 10 units RNaseOUT (Invitrogen; Carlsbad, CA, USA) for 1 hr at 42°C, 10 minutes 50°C, 15 minutes 70°C, hold at 4°C and store at -20°C overnight.

The first round of PCR was performed using 1X Phusion High-Fidelity PCR Master Mix with HF Buffer (New England BioLabs; Ipswitch, MA, USA) with forward primer “reverse transcription primer (RTP)”, and reverse primer “lncCOBRA1 5’ RACE Primer 1” with cDNA diluted 1:5 with the following program: 95°C for 5 minutes; 30 cycles of 95°C for 30 sec, 55°C for 30 sec, 72°C for 1 minute; 72°C 5 minutes, hold at 4°C. PCR 2 was performed similarly, but with PCR reaction 1 diluted 1:20 as the template and “Internal RA5 Primer” forward primer and either (A) “lncCOBRA1 qPCR Reverse set 1” or (B) “lncCOBRA1 5’ RACE Primer 2” as the reverse primer. The PCR reaction was then run on a 1% agarose TAE gel with a 1 kb plus DNA ladder (Invitrogen; Carlsbad, CA, USA). All primers are listed in **Supplemental Data Set 2**.

#### 3’ RACE

To ligate the 3’ adapter, the 3’ RNA adapter (RA3; 5’-TGGAATTCTCGGGTGCCAAGG - 3) was first heated to 70°C for 3 minutes and snapped cool on ice for 2 minutes. 8 μL heat-treated 5 μM RA3 was added to 1 μg RNA isolated from 5-day-old Col-0 seedlings and incubated with 200 units T4 RNA Ligase 2, truncated (New England BioLabs; Ipswitch, MA, USA) in 1X T4 RNA Ligase Buffer (New England BioLabs; Ipswitch, MA, USA) for 1 hr 15 minutes at 28°C. As a control, this reaction was also performed in the absence of T4 RNA Ligase 2, truncated (-Lig). The reaction was then ethanol precipitated overnight.

The following day, the precipitated RNA was split in half for reverse transcription +/- RT. To 8 μL RNA, 1 μL 10 mM dNTPs and 1 μL RTP was added and incubated for 5 minutes at 65°C then transferred to ice for 2 minutes. Reverse transcription was performed with 100 units SuperScriptII (SSII) in 1X First Strand Buffer, 10 mM DTT, 20 units RNaseOUT (Invitrogen; Carlsbad, CA, USA) for 2 minutes at 25°C, 90 minutes 42°C, 5 minutes 95°C, hold at 4°C and store at -20°C overnight. The reaction was also performed without SSII as a control.

The first round of PCR was performed using 1X Phusion High-Fidelity PCR Master Mix with HF Buffer (New England BioLabs; Ipswitch, MA, USA) with cDNA diluted 1:10 in water, and forward primer “RTP” and reverse primer “Illumina RNA index primer 35” with the following program: 95°C for 5 minutes; 30 cycles of 95°C for 30 sec, 55°C for 30 sec, 72°C for 1 minute; 72°C 5 minutes, hold at 4°C. PCR 2 was performed similarly, but with PCR reaction 1 diluted 1:20 as the template and “lncCOBRA1 3’ RACE Primer 1” forward primer and “RNA primer index universal” as the reverse primer. The PCR reaction was then run on a 1% agarose TAE gel with a 1 kb plus DNA ladder (Invitrogen; Carlsbad, CA, USA), excised and gel extracted using the Monarch DNA Gel Extraction Kit following the manufacturers’ instructions (New England BioLabs; Ipswitch, MA, USA).

The purified PCR reaction was then A-tailed with 15 units Klenow Fragment (3’ – 5’ exo-) (New England BioLabs; Ipswitch, MA, USA) in 1X NEB Buffer 2 (New England BioLabs; Ipswitch, MA, USA), and 0.1 mM dATP for 30 minutes at 37°C. The reaction was then cleaned up using Zymo ChIP DNA Clean & Concentrator following the manufacturers’ instructions (Zymo Research; Irvine, CA, USA). The resulting PCR reaction was then cloned into pGEM T-Vector system and selected for using the XGal/IPTG system (Promega, Madison, WI, USA). Sanger sequencing was performed at the University of Pennsylvania Genomic Analysis Core with the SP6 promoter/primer. All primers are listed in **Supplemental Data Set 2**.

### Denaturing RNA gel

Gel was performed using NorthernMax reagents (Invitrogen; Carlsbad, CA, USA). 10 μg of total RNA for each genotype was added to appropriate amount of 3X NorthernMax Formaldehyde Loading dye, boiled at 65°C for 15 minutes and flash cooled on ice. 0.5 μL of ethidium bromide (10mg/mL) was added to each sample and loaded onto a 1.5% NorthernMax denaturing agarose gel and run for ∼3 hr at 100V. Gel was visualized by UV.

### Germination

For germination experiments, seeds of Col-0, *lnccobra1-1*, *lnccobra1-2*, and *lncCOBRA1/lnccobra1-1* were sterilized in 100% ethanol for one-minute followed by a 10-minute wash with 30% Clorox and 0.01% Tween-20 and washed 5X with sterilized water. Seeds were then plated on ½ MS agar plates with 1% sucrose and 0.8% Phytoblend and stratified by cold treating at 4°C for 48 hrs and placed in growth chambers. Two days after transfer to growth chambers, the number of seeds that that displayed cotyledons entirely emerged from the seed coat were counted. Plates were then allowed to grow for 3 more days and 5-day-old seedlings were collected to measure *lncCOBRA1* abundance.

### Leaf initiation rate

Col-0, *lnccobra1-1*, *lnccobra1-2*, and *lncCOBRA1/lnccobra1-1* were grown in soil as described above. Every day at ∼11 AM the presence of leaf primordia was examined. Leaf initiation was measured when the leaf primordia was visible to the eye (∼0.5 mm). After 3 weeks, plants were weighed for fresh weight measurements. To measure plant size, 3-week-old plants were taped flat on paper, scanned, and analyzed using ImageJ as follows. Scanned images were first converted to 8-bit and processed into a binary image such that any plant tissue was converted to white and background became black. Threshold was set using default settings, inverted, and the ‘particles’ (plants) perimeter and area measured. Area of leaf 3 was selected by hard and measured.

### Chromatin Isolation by RNA Purification (ChIRP)

#### Probe design, crosslinking and chromatin isolation

ChIRP probes were designed using the Stellaris probe website (https://www.biosearchtech.com/support/tools/design-software/chirp-probe-designer) with a 3’ Biotin TEG. 5-day-old Col-0 and *lnccobra1-2* seedlings were crosslinked in PBS with 1% formaldehyde (v/v) (Sigma-Aldrich; St. Louis, MO, USA) added and placed under vacuum for 10 minutes, followed by a 5-minute quench with 125 mM Glycine under vacuum. Crosslinked tissue was then washed 5 times in distilled, deionized water, patted dry with paper towels, flash frozen in liquid nitrogen, and stored at -80°C until further processing. Chromatin from 6 g of 5-day-old Col-0 and *lnccobra1-2* crosslinked seedlings (3 g scrambled probes and 3 g *lncCOBRA1* probes) was isolated as previously described (Do et al., 2019). All probes are listed in **Supplemental Data Set 2**.

#### Bead preparation

Pierce High Capacity Streptavadin Agarose beads (Thermo Fisher Scientific; Carlsblad, CA, USA) were first chemically treated to protect the streptavidin from tryptic proteolysis in preparation for mass spectrometry to reduce streptavidin signal as previously described (Barshop et al., 2019).

#### ChIRP

ChIRP was performed as previously described (Chu et al., 2011, 2012, 2015), with several modifications. Modified Pierce High Capacity Streptavadin Agarose beads (Thermo Fisher Scientific; Carlsbad, CA, USA) were first washed twice and resuspended in nuclei lysis buffer (50 mM Tris-HCl pH = 8.0, 10 mM EDTA, 1% SDS) supplemented with cOmplete Protease Inhibitor Cocktail (Sigma, St Louis, MO, USA) and RNaseOUT (Invitrogen, Carlsblad, CA, USA). Chromatin lysates were then pre-cleared with 30 μL modified beads for 30 minutes with mixing in a 37°C hybridization oven with rotation. After pre-clearing, samples were centrifuged twice at 3000 RPM for 5 minutes at room temperature (RT) to thoroughly remove any beads, and 10% of the sample was removed for both RNA input and protein input. The lysates were then split into a scrambled and *lncCOBRA1* probe sample and 2X Hybridization buffer was added (750 mM NaCl, 1% SDS, 50 mM Tris-HCl pH = 7.5, 1 mM EDTA, 15% Formamide) supplemented with PMSF (100 μL/10 mL), RNaseOUT (5 μL/10 mL; Invitrogen, Carlsblad, CA, USA), and cOmplete Protease Inhibitor Cocktail (Sigma, St Louis, MO, USA). 100 pmol of probes were then added per 1 mL chromatin (i.e. 1.67 μL for each of the 6 probes used for *lncCOBRA1*) and the samples incubated in a 37°C hybridization oven with rotation.

After 5 hours, 100 μL of modified beads were added to each tube and incubated in a 37°C hybridization oven with rotation for another 2 hrs. Samples were then centrifuged for 5 minutes at 3000 RPM, supernatant removed, and resuspended in 1 mL wash buffer (2S SSC, 0.5% SDS) pre-warmed to 37°C and incubated in a 37°C hybridization oven with rotation for another 30 minutes. Samples were washed for a total of 4 washes. After the last spin, samples are resuspended in 1 mL wash buffer and 150 μL removed for RNA isolation and the remaining 850 μL used for mass spectrometry.

#### RNA isolation

RNA isolation was performed using a modified version of a previously published protocol (Desvoyes et al., 2018). RNA samples were centrifuged at 3000 RPM for 5 minutes and resuspended in 400 μL RNA proteinase K buffer (PK Buffer; 100 mM NaCl, 10 mM Tris-HCl pH = 7.5, 1mM EDTA, 0.5% SDS) and 390 μL PK buffer was added to RNA input samples. To reverse crosslinks, NaCl was added to a final concentration of 200 mM (add 8 μL 5M NaCl) and incubated at 65°C overnight. The following day 16 μL 1M Tris-HCl pH = 6.8, 8 μL 0.5 M EDTA and 2 μL proteinase K (Denville Scientific; Metuchen, NJ, USA) was added and incubated at 37°C for 2 hr with rotation to remove proteins. Samples were then added to 700 μL Qiazol (Qiagen; Valencia, CA, USA), and RNA isolated as described above. All primers are listed in **Supplemental Data Set 2**.

#### ChIRP-MS qRT-PCR validation

qRT-PCR was performed as described above with the following exceptions. A standard curve for all primer sets was generated using serial dilutions of genomic DNA. “Copy number” of each transcript was calculated, and normalized by the average CT value for three technical replicates of *U6* for each sample. The normalized values were then used to calculate fold enrichment relative to Col-0 input.

#### Mass spectrometry sample preparation and acquisition

Protein samples were centrifuged at 3000 RPM for 5 minutes, supernatant removed, and the beads were wash 3 times with 100 mM NH4HCO3, and ultimately resuspended in 400 μL 100 mM NH4HCO3 supplemented with 200 mM NaCl and incubated overnight at 65°C to reverse crosslinks. The next day the samples were flash frozen in liquid nitrogen and stored at -80°C until processing. Samples were thawed on ice and resuspended in an appropriate volume of the resuspension buffer (50 mM SDS and 50 mM triethylammonium bicarbonate (TEAB) final concentrations) and reduced with final 10 mM DTT (US Biological, Salem, MA, USA) for 30 min at 30 °C, followed by alkylation with final 50 mM iodoacetamide (Sigma, St Louis, MO, USA) for 30 min at 30 °C. The samples were processed using an S-Trap^TM^ column according to the protocol recommended by the supplier (Protifi; Farmingdale, NY,USA; C02-mini): loaded onto the column and digested with trypsin (Thermo Fisher Scientific; Carlsblad, CA, USA) in 1:10 (w/w) enzyme/protein ratio for 1 h at 47 °C.

Peptides eluted from this column were vacuum-dried and resuspended with LC-MS grade water containing 0.1% (v/v) TFA for mass spectrometry analysis. Each sample was analyzed by a Q-Exactive HF mass spectrometer (Thermo Fisher Scientific; Carlsblad, CA, USA) coupled to a Dionex Ultimate 3000 UHPLC system (Thermo Fisher Scientific; Carlsblad, CA, USA) equipped with an in-house made 15 cm long fused silica capillary column (75 μm ID), packed with reversed-phase Repro-Sil Pur C18-AQ 2.4 μm resin (Dr. Maisch; GmbH, Ammerbuch, Germany) column. Elution was performed using a gradient from 5% to 35% B (50 min), followed by 90% B (10 min), and re-equilibration from 90% to 5% B (5 min) with a flow rate of 400 nL/min (mobile phase A: water with 0.1% formic acid; mobile phase B: 80% acetonitrile with 0.1% formic acid). Data were acquired in data-dependent MS/MS mode. Full scan MS settings were: mass range 200−1500 m/z, resolution 120,000; MS1 AGC target 1E6; MS1 Maximum IT 100. MS/MS settings were: resolution 30,000; AGC target 5E4; MS2 Maximum IT 200 ms; fragmentation was enforced by higher-energy collisional dissociation with stepped collision energy of 25, 27, 30; loop count top 15; isolation window 1.4; fixed first mass 120; MS2 Minimum AGC target 2E3; charge exclusion: unassigned, 1, 7, 8 and >8; peptide match preferred; exclude isotope on; dynamic exclusion 45 sec.

#### Mass spectrometry data analysis

The acquired data were processed via Proteome Discoverer 2.4 with the default QExactive Precursor Quant and LFQ MPS with SequestHT and Percolator processing template and Comprehensive Enhanced Annotation LFQ and Precursor Quant consensus template with the following parameters. The spectra match with peptide sequence was performed with SequestHT with contaminants.fasta from MaxQuant and Arabidopsis thaliana fasta 2019.04 release, full tryptic digestion, maximum missed cleavage 3, peptide length between 6 to 144, MS1 mass tolerance 10ppm, MS2 mass tolerance 0.02 Da, dynamic modification with oxidation on methionine, acetylation and methionine loss on protein N-terminal, static modification with carbamidomethyl on cysteine. The protein inference and identification validation was performed with Percolator with a 1% false discovery rate (FDR) cut off. Normalization was performed by total peptide amount and scaling mode was set to on all average. Protein Abundance was peptide summed with the top 3 most abundant peptides for each protein.

Proteins were first filtered such that only proteins with abundance scores in at least two biological replicates of the pulldown with the *lncCOBRA1* probes in Col-0 background were considered. Protein abundance in *lncCOBRA1* pulldown was then normalized by the average protein abundance identified using the scrambled probes (*lncCOBRA1*/scrambled) and only proteins that were enriched with the *lncCOBRA1* probes compared to the scrambled sequence probes were examined further (*COBRA1*/scrambled > 1). Enrichment of Col-0 over *lnccobra1-2* background was then calculated as the log2[Col-0/*lnccobra1-2*] and proteins enriched over 1-fold were classified as *lncCOBRA1*-interacting and used for future analyses. We also examined proteins that were present in at least 2 biological replicates of the *lncCOBRA1* pulldown in Col-0 tissue, but absent from scrambled and in 0 or 1 biological replicate of the *lncCOBRA1* pulldown in the *lnccobra1-2* background. Since no protein abundances were found in the scrambled, a fold enrichment could not be calculated.

### Protein-protein interaction (PPI) network

STRING (https://string-db.org/cgi/input.pl?sessionId=QFOfEIHPOaYj&input_page_show_search=on) was used to generate the protein-protein interaction network with medium stringency and clustered into 5 clusters by the k-means clustering algorithm provided. PPI enrichment was calculated by the STRING program (Szklarczyk et al., 2019).

## Abbreviations

lncRNA: long non-coding RNA
lincRNA: long intergenic non-coding RNA
snoRNA: small nucleolar RNA
nt: nucleotide
RBP: RNA binding protein

---

**Supplemental Figure 1:**
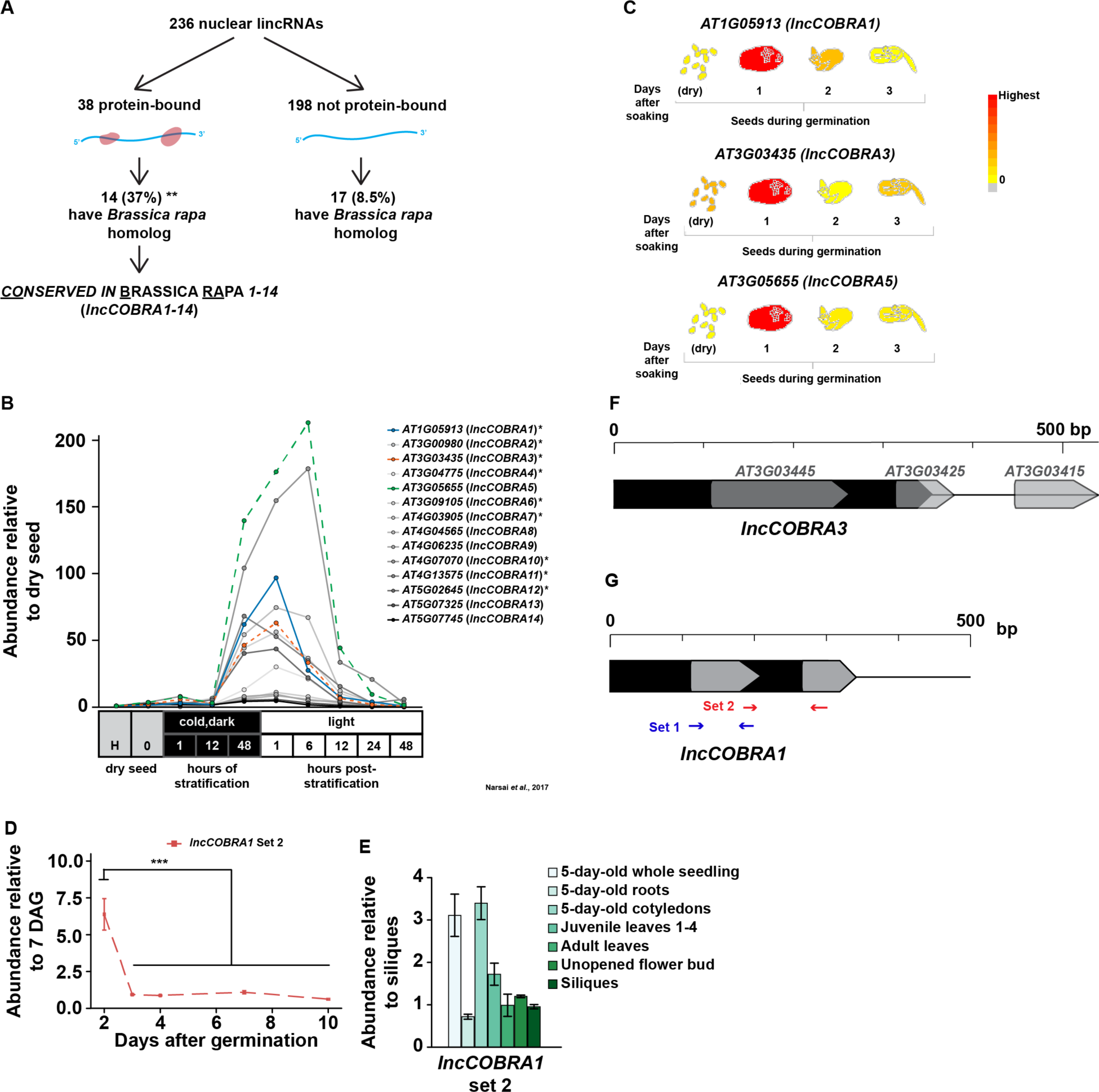
Identification of highly conserved, protein-bound lincRNAs in the nuclei from 10-day-old seedlings. (A) Flowchart diagram of identification of *lncCOBRA* transcripts from protein interaction profile sequencing (PIP-seq) in the nuclei from 10-day-old seedlings (Gosai et al., 2015). (B) Abundance of all *lncCOBRA* transcripts during germination. Abundance is relative to dry seed after harvest. Data was provided in (Narsai et al., 2017). Asterisk denotes lincRNAs with snoRNAs annotated within them. (C) eFP browser views of abundance of *lncCOBRA1*, *lncCOBRA3*, and *lncCOBRA5* during germination (Klepikova et al., 2016). (D) Gene model of *lncCOBRA3* (*AT3G03445*) and nearby snoRNAs. (E) Abundance of *lncCOBRA1* early seedling development using primer set 2 as measured by qRT-PCR. Abundance is normalized by *UBC9* and *UBC10* and is relative to 7-day-old seedlings. *** denotes p-value < 0.001, Wilcoxon t-test. (F) Abundance of *lncCOBRA1* using primer set 2 as measured by qRT-PCR. Abundance is normalized by *UBC9* and *UBC10* and is relative to siliques seedlings. (G) Diagram of *lncCOBRA1* representing the location of the two sets of primers used for qRT-PCR.

**Supplemental Figure 2:**
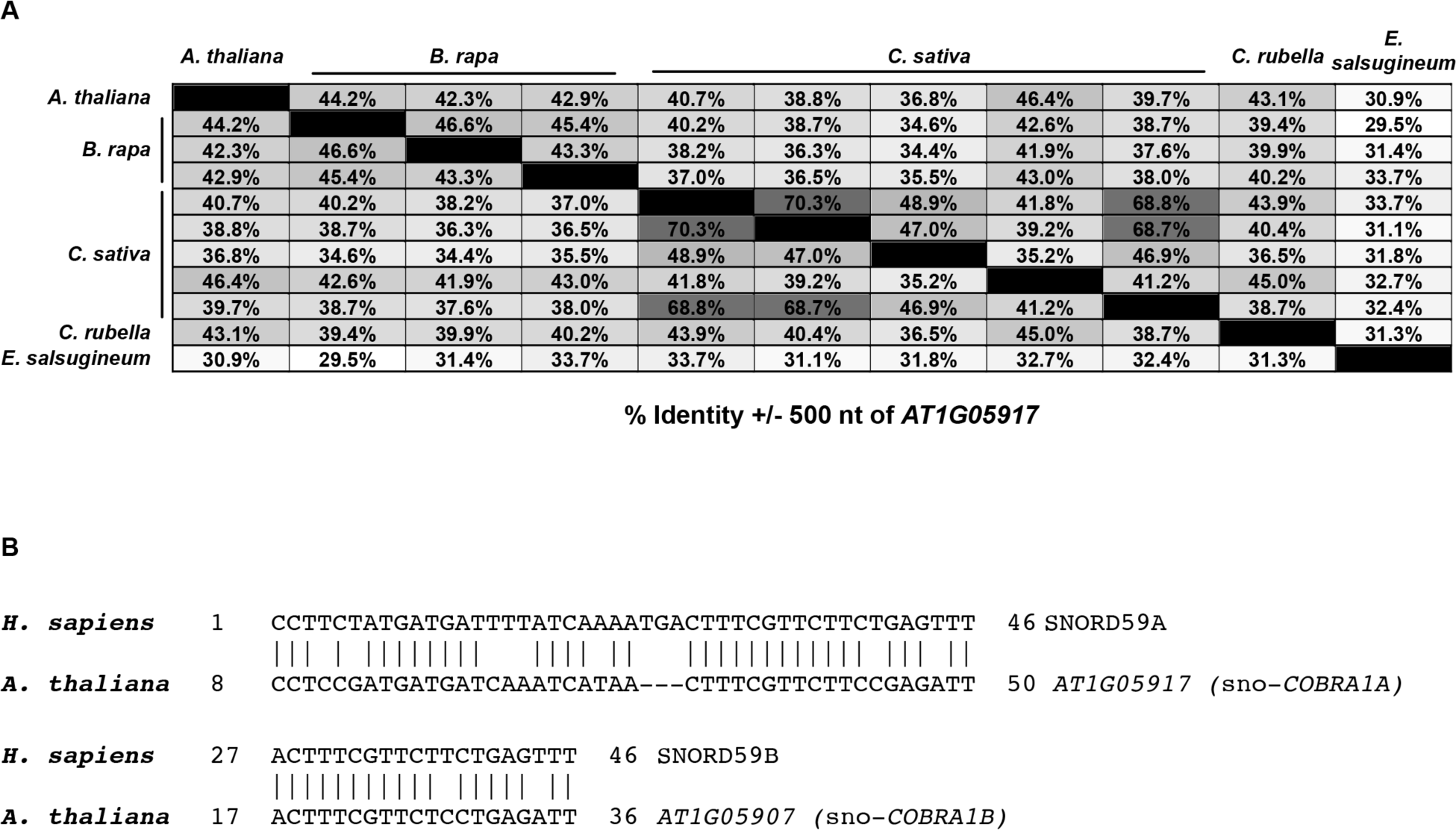
*lncCOBRA1* is highly conserved. (A) Percent nucleotide identity in *Brassica rapa*, *Camelina sativa*, *Capsella rubella*, and *Eutrema salsugineum* in the 500 nt up- and downstream of *AT1G05917* (sno-*COBRA1A*). Calculated by Geneious Prime (Geneious | Bioinformatics Solutions for the Analysis of Molecular Sequence Data, 2019). (B) Comparison between the sequence of sno-*COBRA1A* and sno-*COBRA1B* and their human homologs. Performed using blastn suite from the NCBI aligning two of more sequences (Zhang et al., 2000).

**Supplemental Figure 3:**
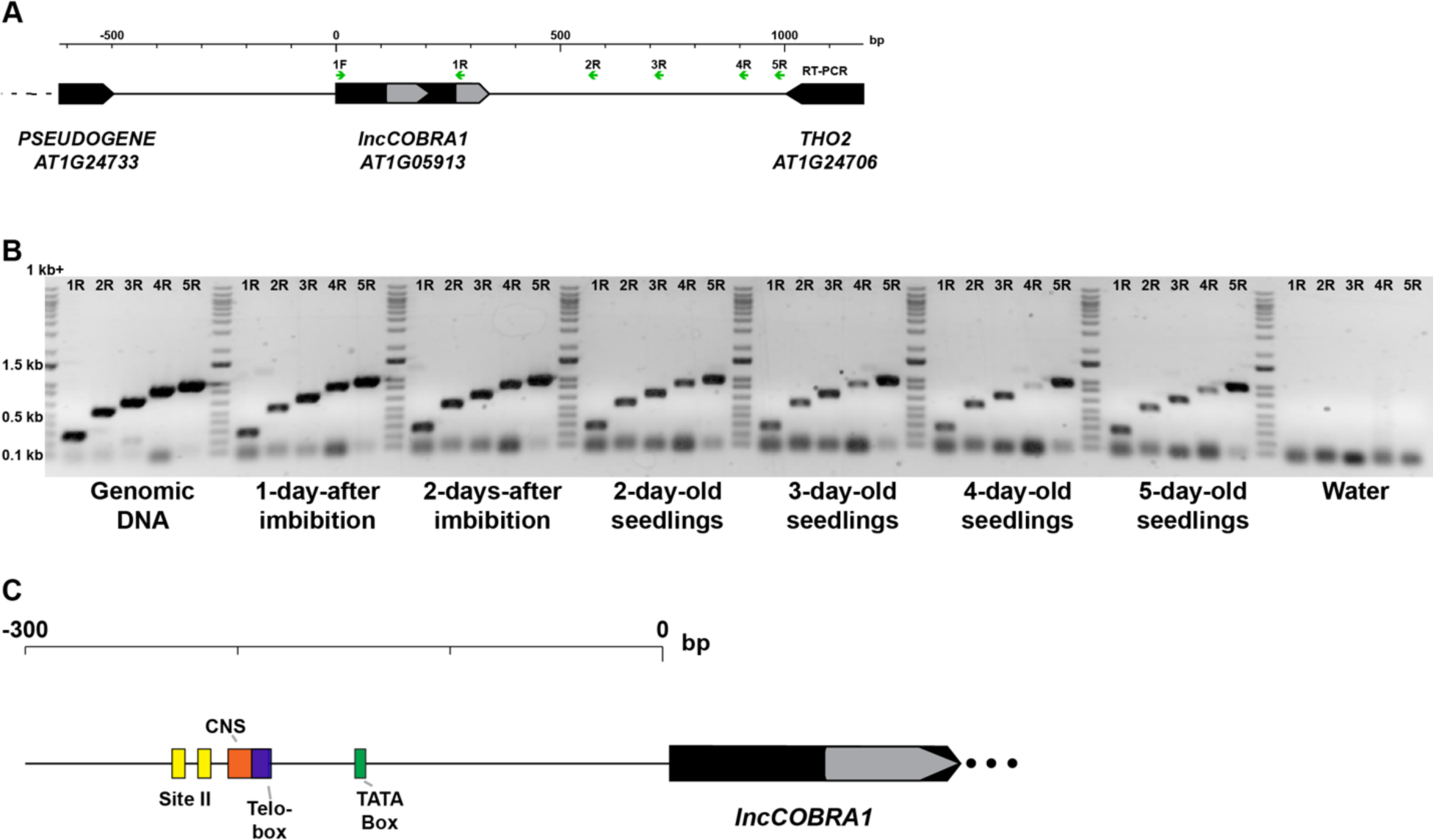
*lncCOBRA1* is transcribed as a longer transcript with conserved promoter elements. (A) Diagram of *lncCOBRA1* representing the location of the RT-PCR primers (green arrows). (B) RT-PCR in cDNA from seeds 1- and 2-day-after soaking in water, and 2-, 3-, 4-, and 5-day-old seedlings. Water is a negative control and genomic Col-0 DNA was used as a positive control. Ladder is 1kb plus. (C) Diagram of the promoter region and conserved elements. Yellow boxes indicate Site II elements, and the purple box represents a Telo-box, together forming a *Telo*SII element. The orange box represents a conserved non-coding sequence (CNS), and the green box represents a TATA-box.

**Supplemental Figure 4:**
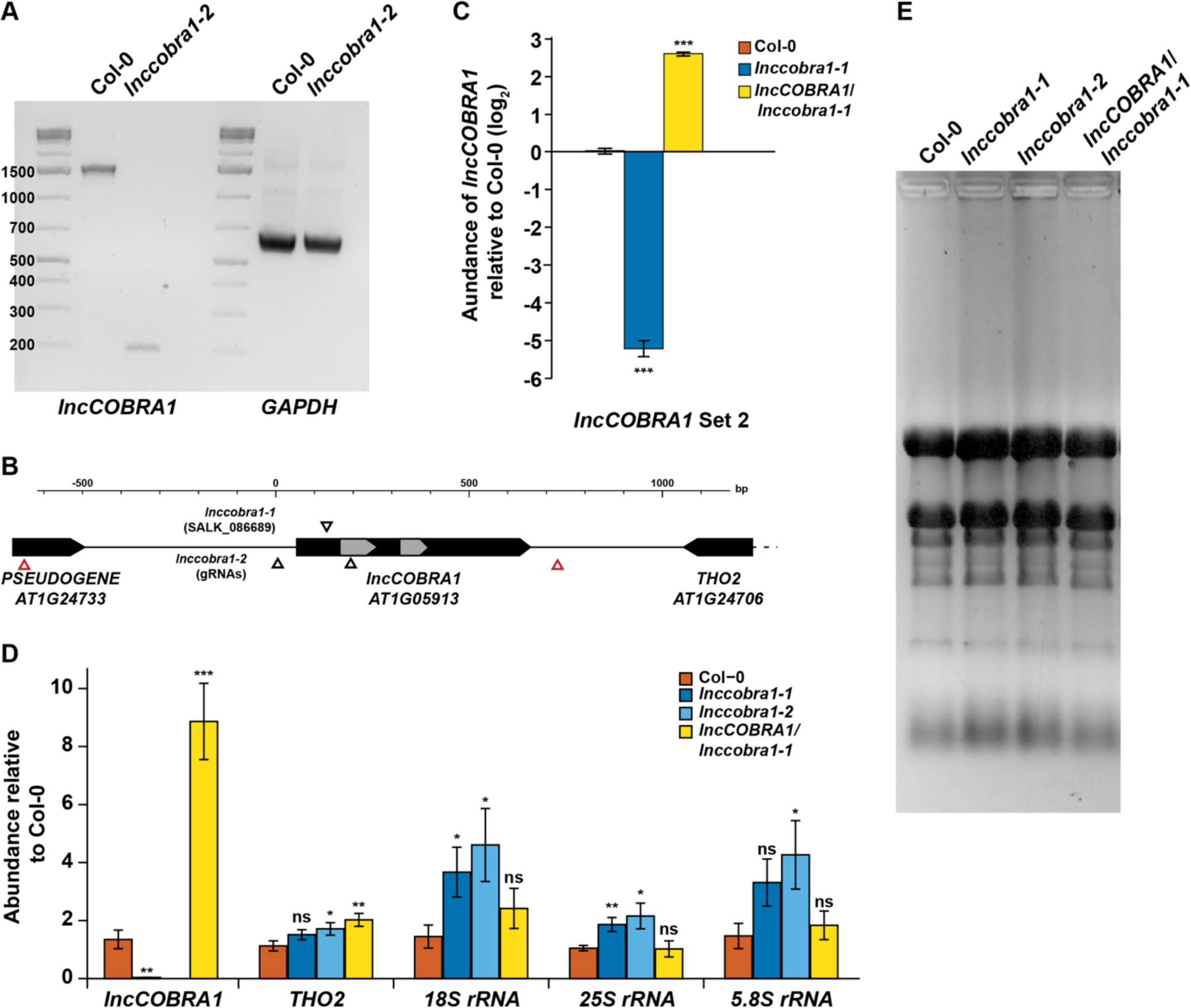
Loss of *lncCOBRA1* minimally affects rRNA abundance. (A) Gel image confirming large deletion caused by two guide RNAs targeted to the 5’ end of *lncCOBRA1*. *GAPDH* is a positive control. (B) Diagram of *lncCOBRA1* representing the location of the T-DNA insertion in *lnccobra1-1*, location of guide RNAs and actual sites of deletion (represented by red triangles) for *lnccobra1-2*. (C) Relative abundance of *lncCOBRA1* in Col-0, *lnccobra1-1,* and *lncCOBRA1*/*lnccobra1-1* using primer set 2 as measured by qRT-PCR. Abundance is normalized by *UBC9* and *UBC10* and is relative to Col-0. *** denotes p-value < 0.001; Wilcoxon t-test. N = 3. Error bars represent SEM. (D) Relative abundance of *lncCOBRA1* (set 1), *5.8S*, *18S*, and *25S rRNA* in Col-0, *lnccobra1-1, lnccobra1-2*, and *lncCOBRA1*/*cobra1-1*. Abundance is normalized by *UBC9* and *UBC10* and is relative to Col-0. *** denotes p-value < 0.001; Wilcoxon t-test. N = 3. Error bars represent SEM. (E) Total RNA isolated from Col-0, *lnccobra1-1, lnccobra1-2*, and *lncCOBRA1/cobra1-1* on a 1.5% denaturing agarose gel.

**Supplemental Figure 5:**
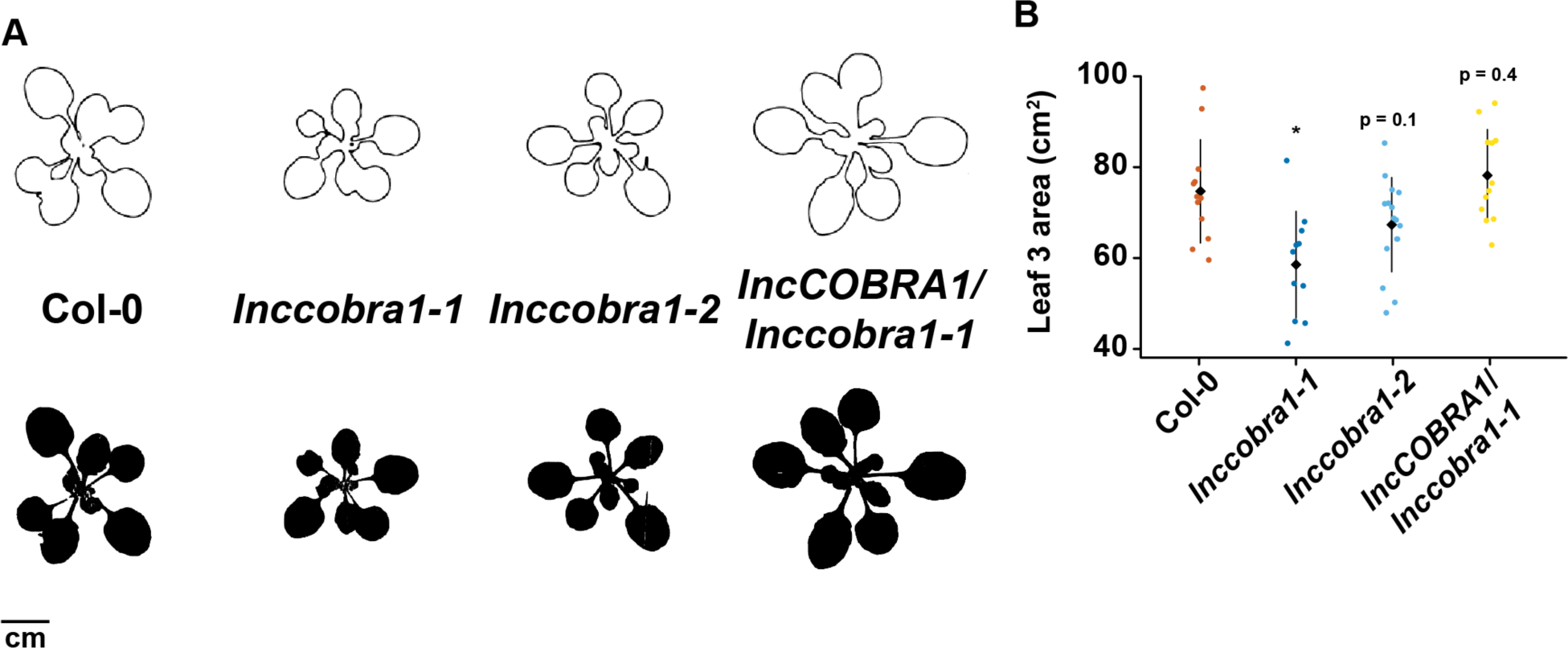
Loss of *lncCOBRA1* results in smaller plants. (A) Representative images generated from ImageJ to measure perimeter of 3-week-old plants. Images on top and bottom are the same plants, with the top being used for perimeter measurements and the bottom used for area measurements. (B) Leaf area of leaf 3 measured by ImageJ. * denotes p-value < 0.05; Wilcoxon t-test.

**Supplemental Figure 6:**
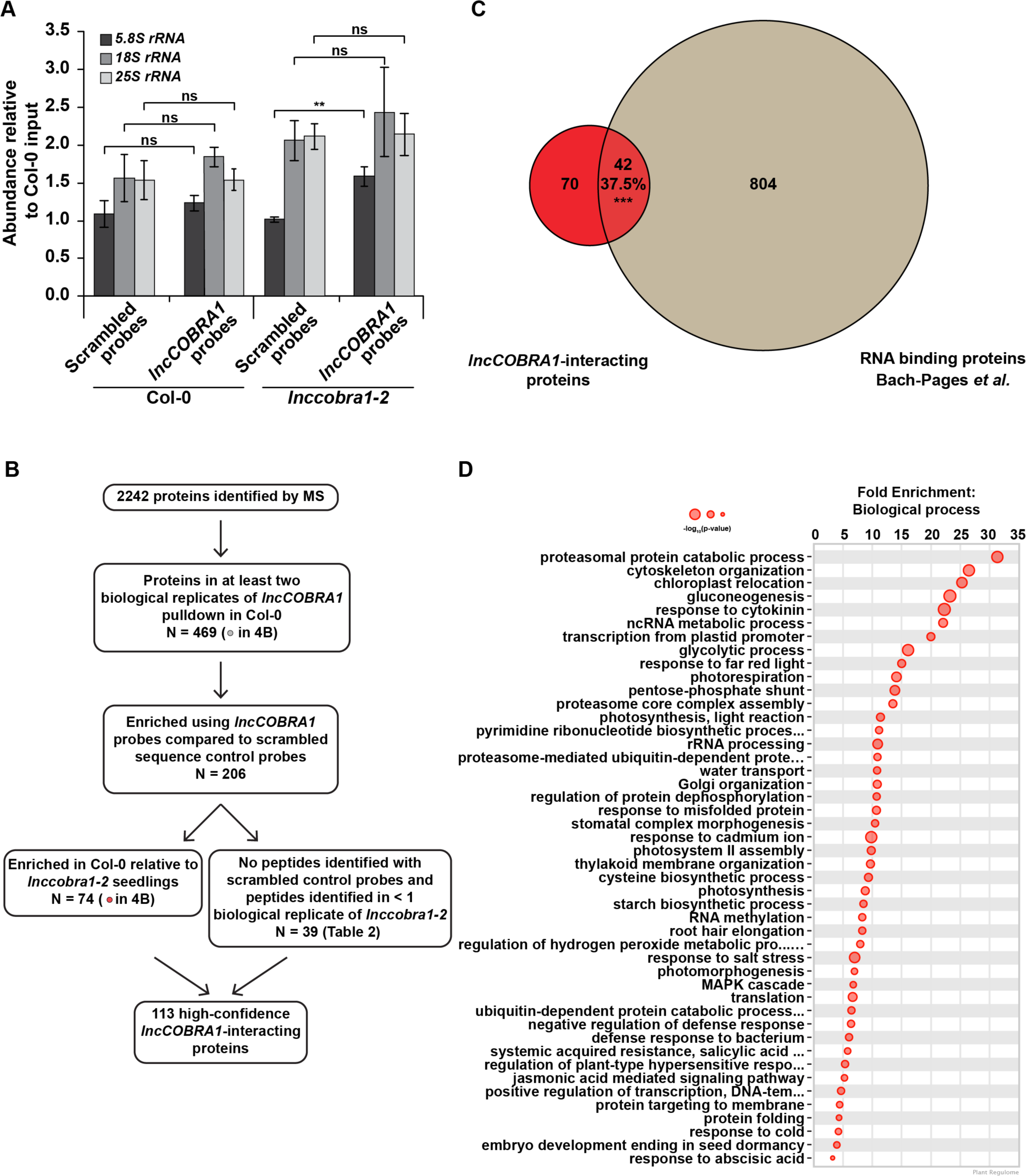
*lncCOBRA1* does not interact with rRNAs. (A) Relative abundance of *5.8S rRNA*, *18S rRNA*, and *25S rRNA* in ChIRP-MS experiments. Abundance is normalized by *U6* and relative to Col-0 input. Error bars represent SEM. ns and **, denotes p-value > 0.05 and < 0.01, respectively; Wilcoxon t-test. N = 3. (B) Overlap between *lncCOBRA1*-interacting proteins and proteins classified as RBPs in an RNA binding proteome capture experiment in Arabidopsis leaves. *** denotes p-value < 0.001; Hypergeometric test. (C) Gene ontology enrichment analysis for biological function using Plant Regulomics (Ran et al., 2020) for *COBRA1*-interacting proteins. Size of circles represents -log_10_(p-value).

**Supplemental Figure 7:**
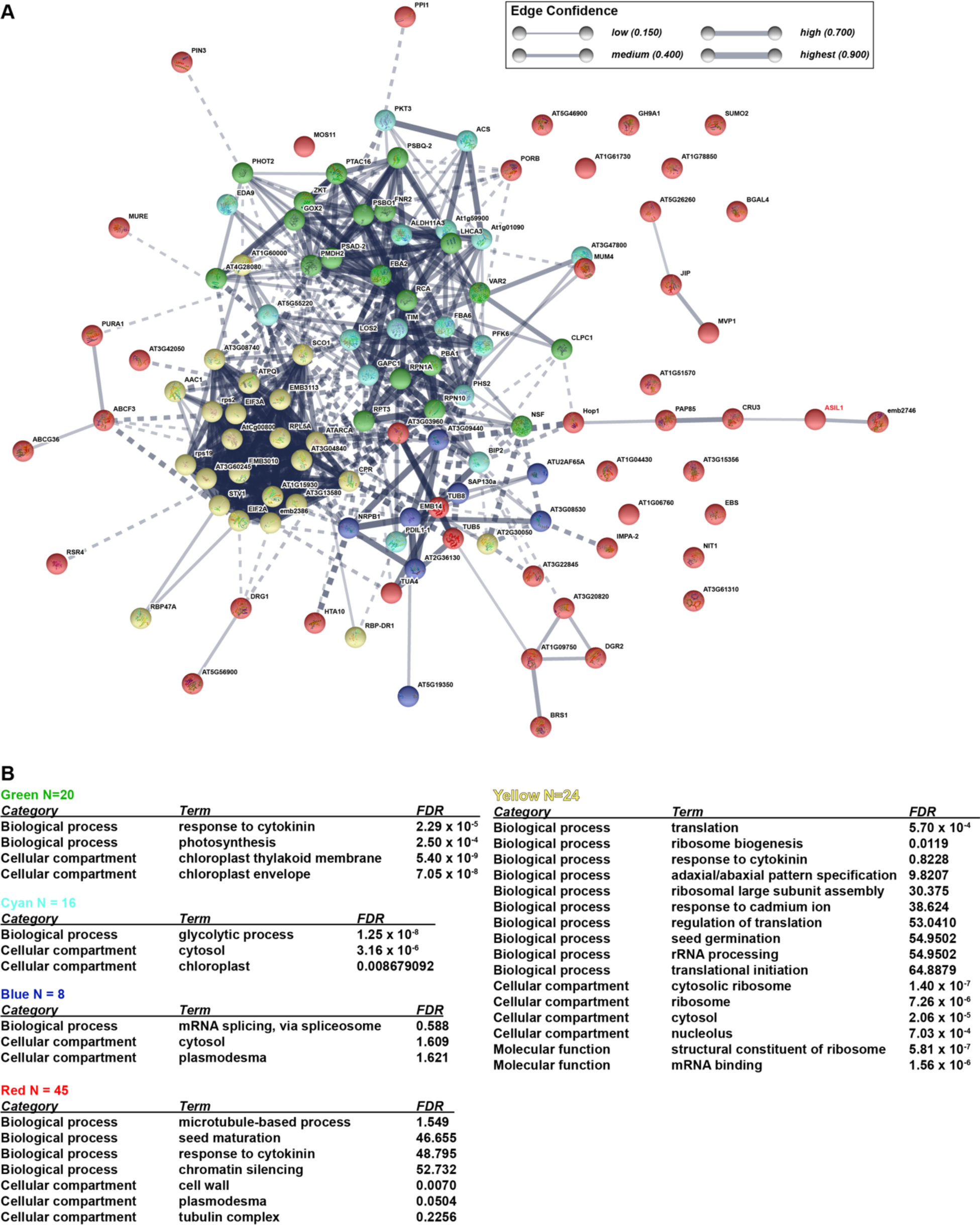
*lncCOBRA1*-interactome. (A) Protein-protein interaction (PPI) network for *lncCOBRA1*-interacting proteins. Proteins were clustered into five clusters by k-means clustering. Thickness of lines connecting notes indicates the confidence of that protein-protein interaction. Dotted line indicates interaction with a different cluster. (B) Gene ontology enrichment for proteins in each cluster (Huang et al., 2009).

## Notes

### Competing Interest Statement

The authors have declared no competing interest.

